# Impact of WaSH and dietary practices on age-driven gut microbiome in stunted young children

**DOI:** 10.64898/2026.03.19.712878

**Authors:** Grantina Modern, Aneth David, Kilaza Samson, Beatus Lyimo, Ianina Altshuler, Sylvester Lyantagaye

## Abstract

**Background:** Stunting, defined as height-for-age below −2 standard deviations of the WHO child growth standards median, is influenced by nutritional and environmental factors. It remains a public health challenge in Tanzania, particularly in Iringa (prevalence 57%, exceeding the national average of 30%), despite abundant food production. This study explored the gut bacteriome as a potential biomarker for child growth and its association with water, sanitation, and hygiene (WaSH) practices in food-secure settings.

**Methods:** A community-based cross-sectional study (September-October 2024) enrolled children aged 5–23 months in Iringa, collecting fecal samples and household data on growth metrics, WaSH, feeding practices, and illness. The V3–V4 region of the 16S rRNA gene was sequenced using Illumina MiSeq and analysed with QIIME2 and R for alpha and beta diversity, differential abundance (ANCOM-BC), and random forest (RF) modelling.

**Results:** Overall, 60.5% of 297 children were stunted. Stunting was associated with older age, male gender, discontinued breastfeeding, poor feeding diversity, toilet sharing, and residence location (p < 0.001, p = 0.049, p = 0.001, p = 0.001, p = 0.001, and p = 0.005, respectively). Significant differences in bacterial community composition were observed between stunted and normally growing children (Shannon p = 0.0053; Bray-Curtis p = 0.001). A shared core bacteriome was identified in both groups, influenced by environmental and dietary factors. Normally growing children were enriched with *Bifidobacterium*, *Rothia*, *Olsenella*, *Slackia*, *Lactobacillus*, *Gemella*, and *Oscillibacter*, while stunted children showed enrichment of *Prevotella*, *Akkermansia*, *Fusobacterium*, *Acinetobacter*, *Alistipes*, *Odoribacter*, *Fournierella*, and the *Ruminococcus torques* group.

**Conclusion:** Age was the most consistent predictor of gut microbial diversity. Stunting does not appear to be caused by a completely different gut microbiome; instead, shared environmental and dietary factors shape both gut bacteria and child growth. Promoting diverse complementary feeding, continued breastfeeding, and improved hygiene could mitigate risks and inform targeted interventions in food-secure regions.

## 1.0 Background

Childhood stunting is defined as a height/length-for-age that is below −2 standard deviations of the WHO child growth standards median (1). Stunting leads to poor physical and cognitive development in children, further causing poor health, education outcomes, and long-term poverty. The WHO aimed to achieve a 40% reduction in the number of stunted children under five by 2025, lowering the global burden from approximately 162 million to 100 million children (2). However, the progress has been insufficient and only about 28% of countries are on track to meet the stunting target by 2030 (3).

Stunting prevalence in Tanzania has gradually decreased from 48.0% in 1999 to 30.0% in 2022, affecting more than half of all children in Iringa (4). Despite numerous efforts to address undernutrition including stunting in Tanzania, most interventions have focused on food security (5). However, emerging evidence links stunting to poor water, sanitation, and hygiene (WaSH) practices (6). In Iringa, food availability alone doesn’t explain the high burden (57%)(7), since food production and consumption is not a problem in the region. Thus, understanding the role of the environment harboring microbes which influence the gut microbiome is important.

Stunting is influenced by both nutritional factors as well as environmental factors. Similar factors influence gut microbiome, a diverse community of microorganisms including bacteria, archaea, fungi, and viruses. These microbes perform numerous essential functions such as food digestion, protection against pathogen overgrowth by stimulating immune system maturation (8), eliminating exogenous toxins and promoting nutrient absorption (9).

The gut microbiome of a child begins forming at birth and is profoundly shaped by environmental microbiomes from surroundings (10). Transmission occurs through multiple pathways, influenced by mode of delivery, feeding practices, hygiene, and living environment (11). These can introduce both beneficial (commensal or symbiotic) and harmful (pathogenic or opportunistic) that may cause dysbiosis, inflammation, or infections (12). In childhood, infants usually have significant diversity of microbiome whereas the composition becomes more stable at between the ages of 1 and 5 years (13). The major composition of gut microbiome belongs to phyla *Firmicutes* and *Bacteroidetes* (14) whereas healthy guts have been associated with abundance of the bacteria phyla *Firmicutes, Bacteroidetes*, and *Proteobacteria* (15).

For children below two years of age, which is a crucial window for growth, WaSH factors such as access to clean water, sanitation facilities, and hygiene practices, significantly influence the composition and diversity of the gut microbiome (16,17). Improved WaSH practices can foster a healthier gut microbiome by reducing pathogenic exposure and supporting beneficial microbial communities, which are essential for digestion, immunity, and growth. Whereas, poor WaSH conditions can expose children to harmful microbes and environmental contaminants, disrupting microbial balance and increasing the risk of diseases like diarrhea, chronic inflammation and malnutrition (18) (19) (20).

Contaminated water or inadequate sanitation, increase exposure to pathogenic bacteria such as *Escherichia coli* or *Salmonella*, which disrupt the gut microbiome by outcompeting beneficial microbes, reducing microbial diversity, cause WaSH-related infections such as diarrhea and triggering inflammation. Also, food prepared in unhygienic conditions can introduce toxins or pathogens such as *Bacillus cereus*, *Campylobacter jejuni*, *Clostridium botulinum*, *Clostridium perfringens*, *Cronobacter sakazakii*, *Esherichia coli*, *Listeria monocytogenes*, *Salmonella* spp., *Shigella* spp., *Staphylococccus aureus*, *Vibrio* spp. and *Yersinia enterocolitica* that alter gut microbial metabolism (21). These may lead to malnutrition including stunting (22).

Exposure to harmful microbes through contaminated water or poor hygiene can overstimulate the immune system, leading to chronic gut inflammation including environmental enteric dysfunction (EED) (23). This inflammatory state alters the gut environment, reduces nutrients absorption and reduces populations of anti-inflammatory genera like *Faecalibacterium*, which are crucial for maintaining gut barrier integrity (24,25).

In areas with poor WaSH, frequent and/or recurrent infections including diarrhea often may lead to increased use of antibiotics to treat these illnesses, which highly reduce microbial diversity, depleting beneficial taxa and promoting antibiotic-resistant strains in the gut. These practices contribute to absence or reduced abundance of key beneficial microbiomes (26), reduce gut integrity and impair growth. The effect of human-microbiome interaction on the host’s gut physiology is evident (8).

Thus, the aim of this study was to explore the gut bacteriome profiles of Iringa children as a potential biomarker for child growth and identify environmental conditions that favor a healthy gut. Specifically, the study’s objectives include characterizing the gut bacteriome profile across age and populations and evaluating the association between gut bacteriome and the influence of WaSH practices in children below 2 years of age.

## 2.0 Methods

### 2.1 Study area

The study was conducted in the Iringa region, located within the Southern Highlands zone of Tanzania mainland: one of eight national geographical zones (Northern, Lake, Central, Western, Eastern, Southern, Southwest, and Southern Highlands). Iringa is administratively encompassed by Iringa urban, Iringa district, Kilolo district, Mafinga district, and Mufindi district. Centered at 7°51′00″S, 35°33′00″E with an average altitude of ∼1,600m above sea level, the region spans approximately 58,370 km² and supports a population of 1,192,728 according to the 2022 preliminary census report, yielding a density of ∼34 persons/km² under a subtropical highland climate. This high burdened, high-altitude setting is characterized by diverse topography and agricultural systems including tea, maize, livestock.

### 2.2 Study design

In September to October 2024, a cross-sectional community-based survey was carried out to understand the reason for high prevalence of stunting despite the region producing adequate food.

### 2.3 Sample size and sampling approach

The study site (Iringa) was stratified based on its districts and wards i.e. Iringa District with 18 wards, Iringa Municipal with 28 wards, Mafinga Town with 9 wards, Mufindi District with 27 wards and Kilolo District with 24 wards. Five out of 106 wards in four districts of Iringa municipal, Iringa district, Mafinga town municipal and Kilolo district were randomly selected based on their population sizes by using ENA for SMART randomization software (UNICEF, 2012). The number of selected children was calculated based on the population size of the selected wards i.e. number of children per ward = ward population/total selected wards population * sample size. From each ward, three (3) villages were conveniently selected based on their distance from the Iringa Regional Referral Hospital in kilometers and the estimated arrival time (ETA) where all samples were stored during the field work. The ward sample size was equally split into 3 to determine the sample size per village.

Fecal samples from children from the 15 villages were collected. A household survey provided metadata including demographic data, child growth measurements (height and weight), WaSH practices, dietary diversity and child illness.

### 2.4 Study population, recruitment, and inclusion/exclusion criteria

Children aged 5 to 23 months were enrolled from selected households across Iringa. Eligible children were required to: (i) fall within the age range, below two years of age; (ii) reside permanently in a study household; (iii) have written informed consent provided by a parent or guardian; (iv) have no self-reported antibiotic use (prescribed or self-medicated) in the preceding 4 weeks; and (v) present without acute or chronic illness or related symptoms at the time of enrollment. Children were excluded if they: (i) were above 24 months of age; (ii) lacked parental consent or withdrew participation; or (iv) had received antibiotics within the prior 4 weeks. Gender and ethnicity were not considered in selection to ensure broad representativeness of the regional infant population.

### 2.5 Sample collection, transportation and storage

Fecal samples were randomly collected from children’s residences by parents or guardians using sterile stool universal containers, with assistance from field assistants and community health workers (CHWs). The samples were then temporarily stored in cooler boxes with ice packs at households and subsequently transferred to a vehicle’s portable freezer (−20°C) for storage during field hours and later transported to the field laboratory at Iringa Regional Referral Hospital. Each sample was thoroughly mixed and aliquoted into 2 mL Eppendorf tubes by laboratory assistant and all aliquots were temporarily stored in a −80°C freezer at the hospital laboratory prior to transportation to the main laboratory for analysis. Upon arrival at the Nelson Mandela African Institution of Science and Technology (NM-AIST) molecular biology laboratory, all aliquots were stored at −80°C pending DNA extraction and genomic sequencing.

### 2.6 Data source

Data for this study were collected through a semi-structured questionnaire and fecal sample analysis. The questionnaire was administered via oral interviews with parents or guardians of the enrolled children to capture socio-demographic characteristics, anthropometric measurements, water, sanitation, and hygiene (WaSH) practices, feeding habits, vaccination history, antibiotic use, and recent illness episodes. Quantitative responses were recorded using digital Kobo Toolbox forms installed on tablets. Subsequently, sequencing data from analyzed fecal samples were documented as paired-end FASTQ files for downstream microbiome profiling and analysis.

### 2.7 Study variables

The primary outcome variable was stunting status, defined as a binary variable where children with height-for-age z-scores (HAZ) <-2 were classified as stunted, and those with HAZ ≥ −2 as normal. Independent variables included socio-demographic and environmental characteristics such as child’s age and gender, childbirth method, mother’s marital status, education and employment, household water sources, distance to water point, sanitation type, toilet sharing, waste disposal practices, and hygiene behaviors (handwashing practices). Child health and nutrition variables entailed illness history, antibiotic use, exclusive breastfeeding duration, extended breastfeeding, vaccination and vitamin A supplementation status, and dietary diversity indicators.

Microbiome-related variables were derived from 16S rRNA sequencing and included alpha diversity indices (shannon diversity and observed features) and beta diversity measures (Bray–Curtis dissimilarity, and Jaccard distance). Taxonomic profiles at phylum and genus levels and relative abundances of dominant taxa were also considered as explanatory variables in relation to stunting, nutrition and environmental factors.

### 2.8 Laboratory analysis

#### DNA extraction

In the laboratory, fecal DNA was extracted using the QIAamp Fast DNA Stool Mini Kit (cat no.51604) (27) at NM-AIST Molecular laboratory as per protocol. The quality and concentration of the DNA were checked through gel electrophoresis and Nanodrop spectrophotometer prior to sequencing.

#### Sequencing

Library preparation followed the Illumina 16S Metagenomic Sequencing Library Preparation Protocol (Part #15044223, Rev. B). The hypervariable V3–V4 region of the bacterial 16S rRNA gene was amplified using universal primers 341F (5′-CCTACGGGNGGCWGCAG-3′) and 805R (5′-GACTACHVGGGTATCTAATCC-3′). PCR amplification was performed using Herculase II Fusion DNA Polymerase with dual indexing via the Nextera XT Index Kit v2. Sequencing was carried out on the Illumina MiSeq platform using 2 × 301 bp paired-end chemistry. All but one of the samples (n = 296) passed library quality control and were successfully sequenced.

### 2.9 Bioinformatics analysis

Bioinformatics data analysis was conducted using the QIIME 2 (version 2025.7) suite of packages (28). Briefly, raw paired-end sequence data, together with the corresponding manifest file, were imported into QIIME 2. The reads were demultiplexed before proceeding with quality check. The resulting summary table and interactive quality plots were used to assess read quality and guide the selection of parameters for denoising. Reads exhibited high quality, ∼91% of the reads had an average phred score of Q20 > 80% with GC content ranging from 50–58%.

DADA2 was used for quality control, where low-quality sequences and chimeras were removed. Trimming was done for both forward and reverse reads by removing10 bp, while truncating was done at 260 bp and 240 bp for forward and reverse reads, respectively. Final read counts per sample exceeded 10,000 reads for all but two. Taxonomic assignment of amplicon sequence variants (ASVs) from quality control was performed using a pre-trained region-specific GSR-DB V3–V4 classifier (29). To evaluate the association between the gut bacteriome and WaSH practices, as well as other environmental factors. Statistical analyses were conducted in R version 4.5.1 (30), where the QIIME2 output ASV table, representative sequences and taxonomy table/file were imported for further downstream statistical analysis.

#### Gut microbial diversity

Alpha diversity indices i.e. shannon and observed features were grouped analysed for variables such as stunting, toilet sharing, handwashing behavior, continued breastfeeding, and feeding practices to show the microbial richness and diversity. Beta diversity was assessed using Jaccard index, and Bray–Curtis dissimilarity, with principal coordinates analysis (PCoA) visualized through Emperor plots with statistical significant confirmation by permutational multivariate analysis of variance (PERMANOVA). To evaluate the relationship between children’s age and gut microbial composition, a mantel test was performed with beta diversity distance metrics.

#### Taxonomic composition and differential abundance

Relative abundance was calculated by normalizing genus-level counts to the total reads per sample, and taxa were ranked based on mean relative abundance across all samples. Relative abundance entailed the common genera at top 30 and phyla at top 10 abundant taxa based on the variable. Core gut bacterial genera, defined as taxa present in more than 50% of samples within study population or within each group, were identified with a prevalence threshold of ≥50% and a minimum mean relative abundance of 0.1% across the study population and stratified by stunting status to identify the core gut bacteriome.

Differential abundance tested with ANCOM-BC identified taxa associated with stunting and other exposures. Log-fold change estimates indicated the direction of enrichment in each comparison, and taxa were considered significantly different at a false discovery rate (FDR)-adjusted q-value < 0.05. Results were visualized using volcano and bar plots to highlight key genera enriched in each group. To further explore microbial taxa associated with stunting, a random forest (RF) (31) classification model was applied using genus-level relative abundance profiles. The model was trained using 500 trees. Feature importance was evaluated by Mean Decrease in Gini index, reflecting the contribution of each genus to classification accuracy.

### 2.10 Ethical statement

The study’s protocol was approved by the Northern Tanzania Health Research Ethics Committee (KNCHREC) with approval number KNCHREC00021/01/2024 renewed to KNCHREC00021/01/2024/b. Written informed consent was obtained from parents/guardians of the selected children.

### 2.11 Declaration for AI tool usage

The authors used Claude (Anthropic), a large language model, solely for grammar checking and language polishing of the manuscript. No AI-generated content was incorporated into the scientific findings, data analysis, interpretation, or conclusions. All intellectual contributions, including study design, data collection, analysis, and writing of the original manuscript, were performed exclusively by the authors.

## 3.0 Results

### 3.1 Study population characteristics

A total of 297 children aged 5–23 months were enrolled in this study, comprising 51% male and 49% female, with a mean age of 14.7 ± 5.2 months. A higher proportion of male children (boys) was observed among the stunted group (57%) compared with the normal growing children (45%; p = 0.049). Nutritional assessment indicated that stunting affected 60.5% of children, while the overall prevalence of underweight and wasting was 12% and 6%, respectively. The majority of births were through vaginal delivery (70%). Detailed population characteristics are shown in table 1. Statistically, young boys had almost double the odds of being stunted than girls (OR = 1.9, p = 0.011) and children with underweight had 6 times higher risks (OR = 6.3, p = 0.0001) (supplementary table 1).

**Table 1:** Population demographics characteristics.

| Characteristic | Normal (N=117)<sup>1</sup> | Stunted (N=179)<sup>1</sup> | P-value<sup>2</sup> |
| --- | --- | --- | --- |
| Child age (months) | 13.1 (5.2) | 15.7 (5.0) | <0.001 |
| Gender |  |  | 0.049 |
| Female | 64 (55%) | 77 (43%) |  |
| Male | 53 (45%) | 102 (57%) |  |
| Delivery mode |  |  | 0.6 |
| Cesarian section | 39 (33%) | 54 (30%) |  |
| Vaginal delivery | 78 (67%) | 125 (70%) |  |
| Household toilet sharing | 64 (55%) | 64 (36%) | 0.001 |
| Extended breastfeeding | 95 (81%) | 114 (64%) | 0.001 |
| Recent child illness | 31 (26%) | 47 (26%) | >0.9 |
| Handwash before feeding | 100 (85%) | 161 (90%) | 0.2 |
| Handwash after feeding | 92 (79%) | 155 (87%) | 0.072 |
| Dietary diversity status |  |  | 0.029 |
| Normal | 106 (91%) | 173 (97%) |  |
| Poor diversity | 11 (9.4%) | 6 (3.4%) |  |
| Geographic residence |  |  | 0.005 |
| Iringa DC | 23 (20%) | 47 (26%) |  |
| Iringa MC | 23 (20%) | 13 (7.3%) |  |
| Kilolo DC | 29 (25%) | 62 (35%) |  |
| Mafinga DC | 42 (36%) | 57 (32%) |  |
| Child handwash before food | 84 (72%) | 154 (86%) | 0.003 |
| Child handwash after food | 74 (63%) | 139 (78%) | 0.007 |

### 3.2 Dietary diversity and WaSH factors

Dietary assessment showed that a small portion of children (13%) had poor feeding diversity. Breastfeeding practices were encouraging; nearly 83% of children were exclusively breastfed for six months, and 72% continued breastfeeding at the time of the survey. Continued breastfeeding was more prevalent among normal growing children (81%) than stunted ones (64%) ( p = 0.001).

In terms of hygiene, 79 % of caregivers reported that the hands of the child were washed before eating and 71% reported handwashing after eating. Household sanitation was generally good, although 46% of households reported sharing toilet facilities, which was more frequent among households with stunted children (36%) than normal ones (55%) (p = 0.001) (table 1).

Geographically, the largest proportions of participants were from Mafinga District (32%), and Kilolo District (35%), then Iringa District Council (26%), and Iringa Municipal Council (7%) (p = 0.005). Children living at Makorongoni ward of Iringa Municipal Council had 20% less chance of being stunted (OR = 0.265, P = 0.03) (table 1).

### 3.3 Gut microbial diversity

#### 3.3.1 Stunting

The alpha diversity in the stunted and normal growing children was analyzed using the Shannon and observed richness indices. This study shows that stunted children had higher microbial diversity compared to children with normal growth (Shannon: p = 0.0053, Observed richness: p = 0.0025). The Shannon index shows good diversity from both groups (H = 3.3, H = 3.5), however the stunted children had richer and more even species (figure 1).

**Figure 1:**
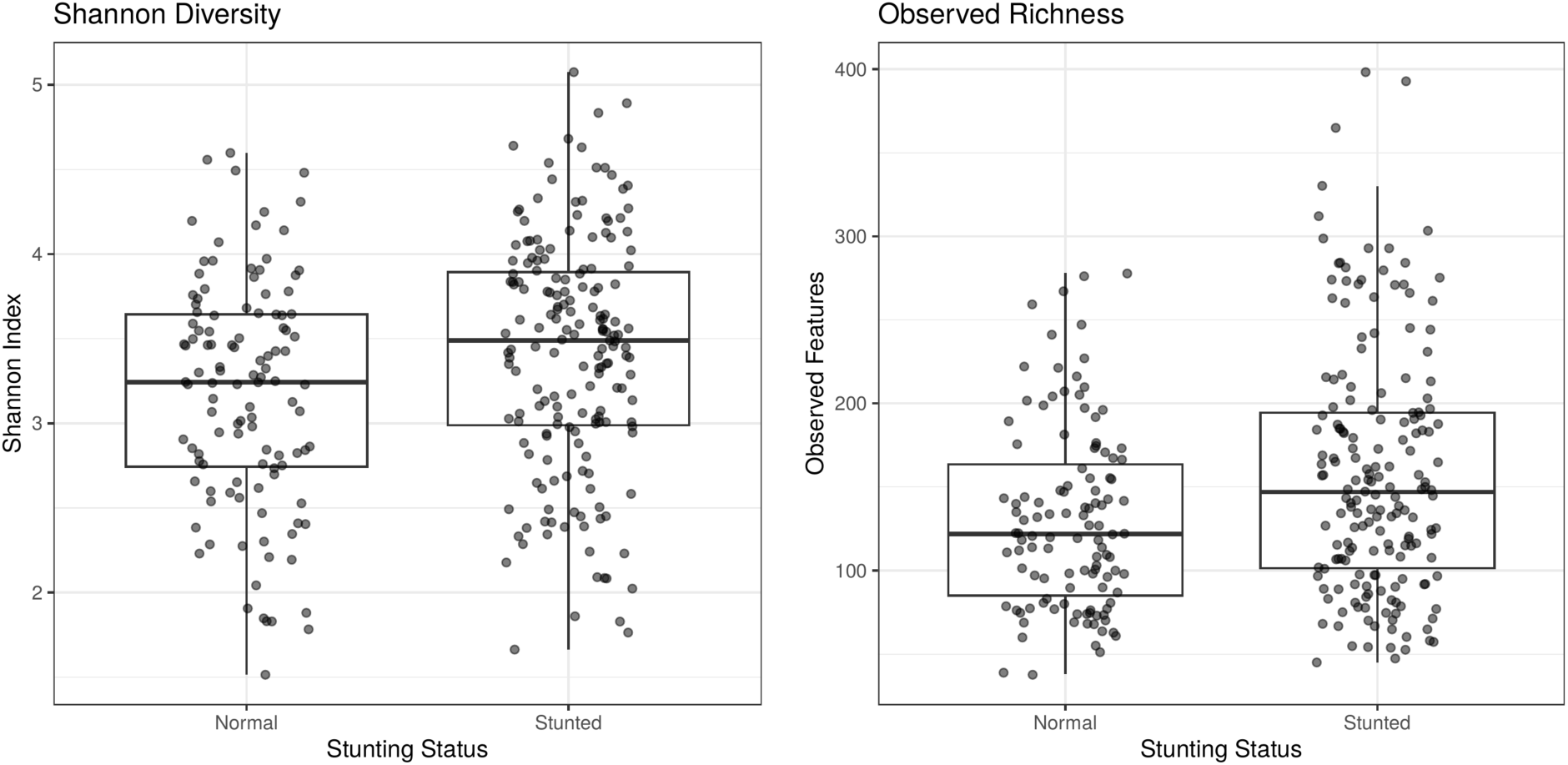
Association between child stunting and gut microbial diversity. Alpha diversity metrics were compared between stunted and normal-growing children. Boxes represent interquartile ranges (IQR), horizontal lines the median, and whiskers 1.5×IQR

While microbial diversity appeared higher in stunted children, they were also significantly older (χ² = 17.28, p = 3.22 × 10⁻⁵). A statistical correlation test shows that age is the most consistent predictor of gut microbial diversity across alpha metrics (rho: 0.51, p < 2.2e-16) (Figure 2).

**Figure 2:**
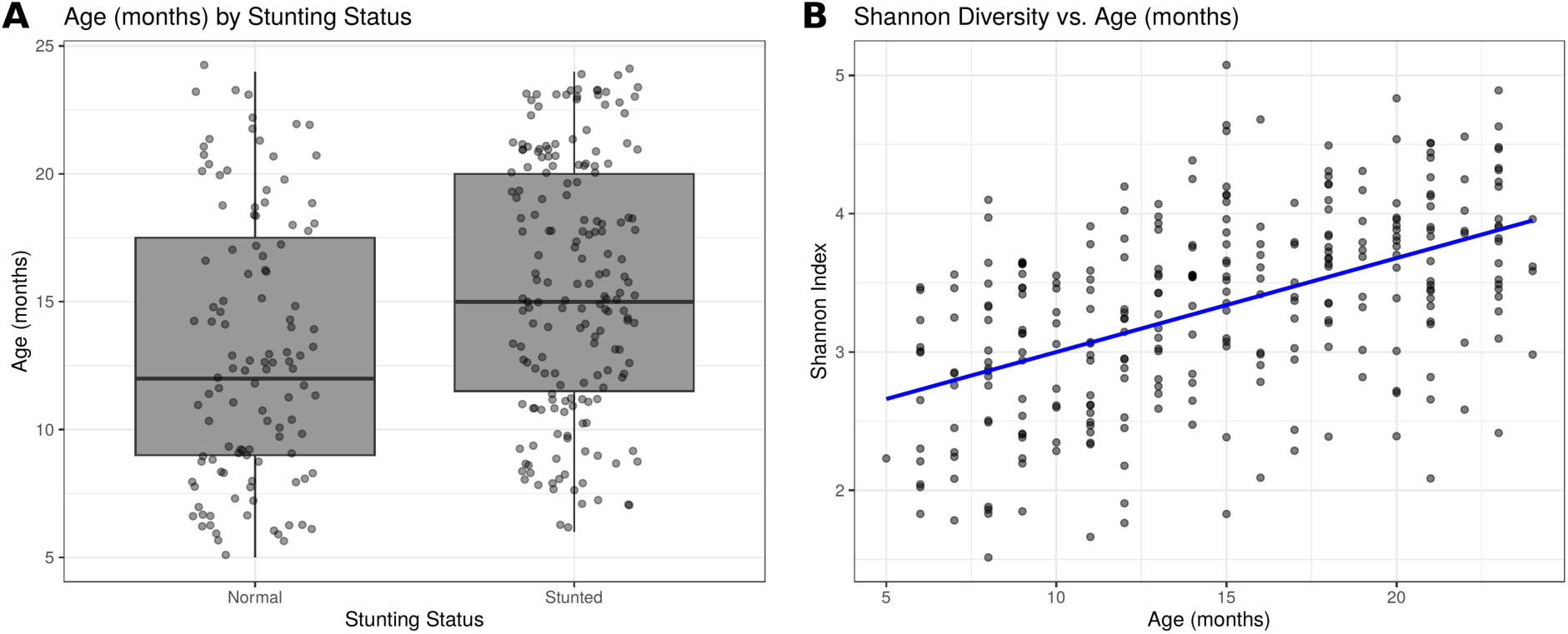
Association between child stunting and age in months. Figure 2A shows the bar plot for stunting status and age. Figure 2B shows the mantel test correlation between Shannon index for stunting status and age in months. Boxes represent interquartile ranges (IQR), horizontal lines the median, and whiskers 1.5×IQR.

Beta diversity through Bray-curtis and Jaccard was used to describe gut microbiota variation between groups including the stunted and normal growing children. Figure 3A shows that microbial composition of stunted children differed significantly from that of normal children. PCoA plots show no clear separation between the children, however there is a subtle statistical difference (p = 0.001; p = 0.001).

**Figure 3:**
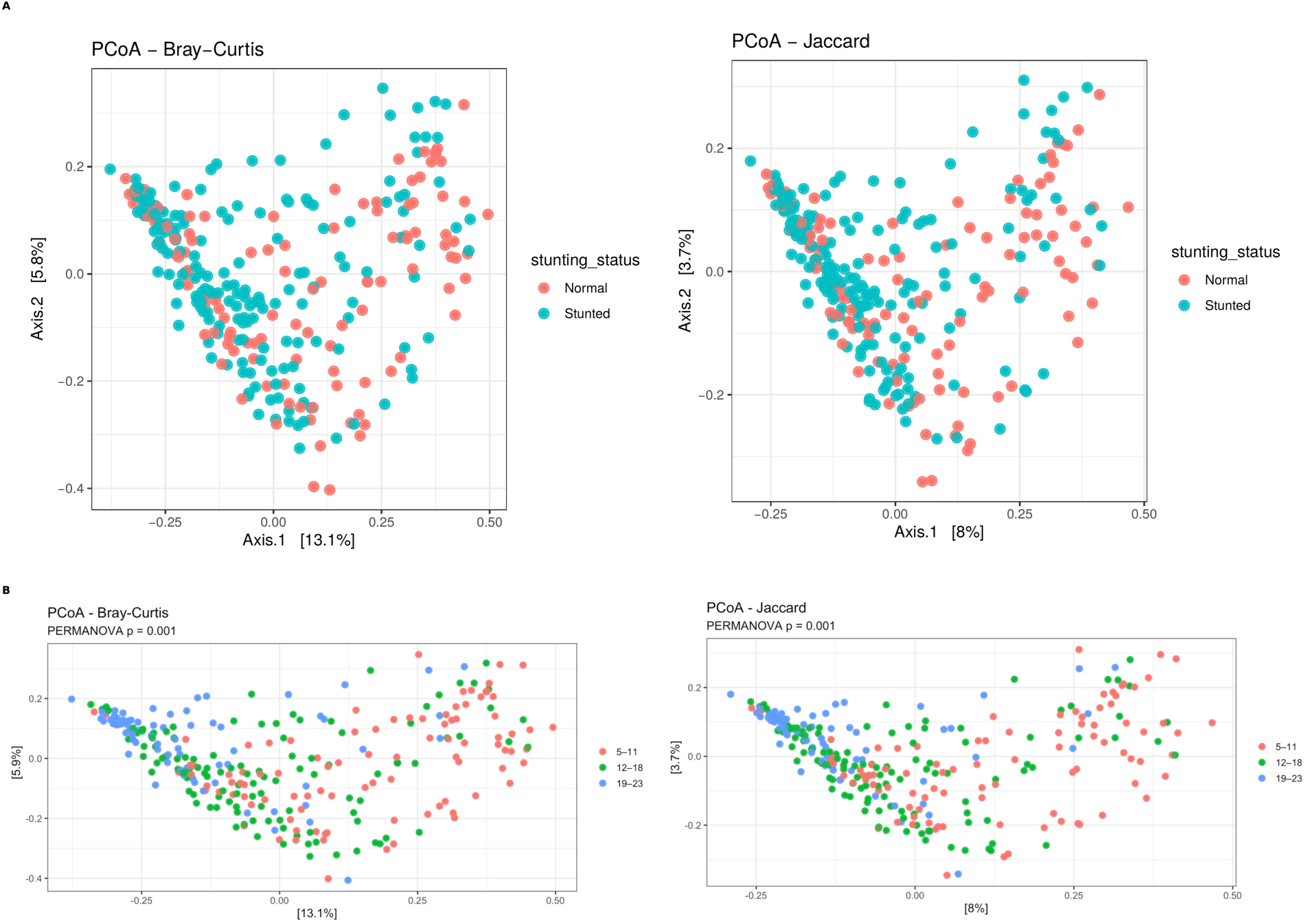
Beta diversity of gut microbial communities by stunting status and age. Figure 3A shows the Principal Coordinates Analysis (PCoA) plots based on Bray–Curtis, and Jaccard illustrate gut microbial communities between stunted and normal-growth children. The Bray-Curtis axes explained 13.1% and 5.8% of the total variation in microbial community composition, respectively. Figure 3B shows beta diversity of gut microbial communities by age categories. The ordinations stratified by age categories show a slow shift in microbial composition with increasing age.

In addition, when the children were stratified into three age categories i.e. 5-11, 12-18, and 19-23 months, no clear clustering was observed, although older children above 11 months were observed to have a slight different composition to younger infants with a subtle significant (figure 3B).

#### 3.3.2 Covariate influencers

Since the diversity of the gut microbiota can be influenced by various factors, in this study, a further analysis was conducted to determine whether nutritional and environmental factors had effects on the microbial diversity.

The results revealed that children with adequate and recommended dietary diversity (p = 1.1×10⁻⁴), underweight children (p = 0.004), those who were no longer breastfeeding (p < 0.001), and children who were washed hands by their caregivers before or after feeding them (p =0.0001; 0.00003) exhibited higher diversity. Lower microbial diversity was observed in sick children with diarrhea (p = 0.0094) (figure 4).

**Figure 4:**
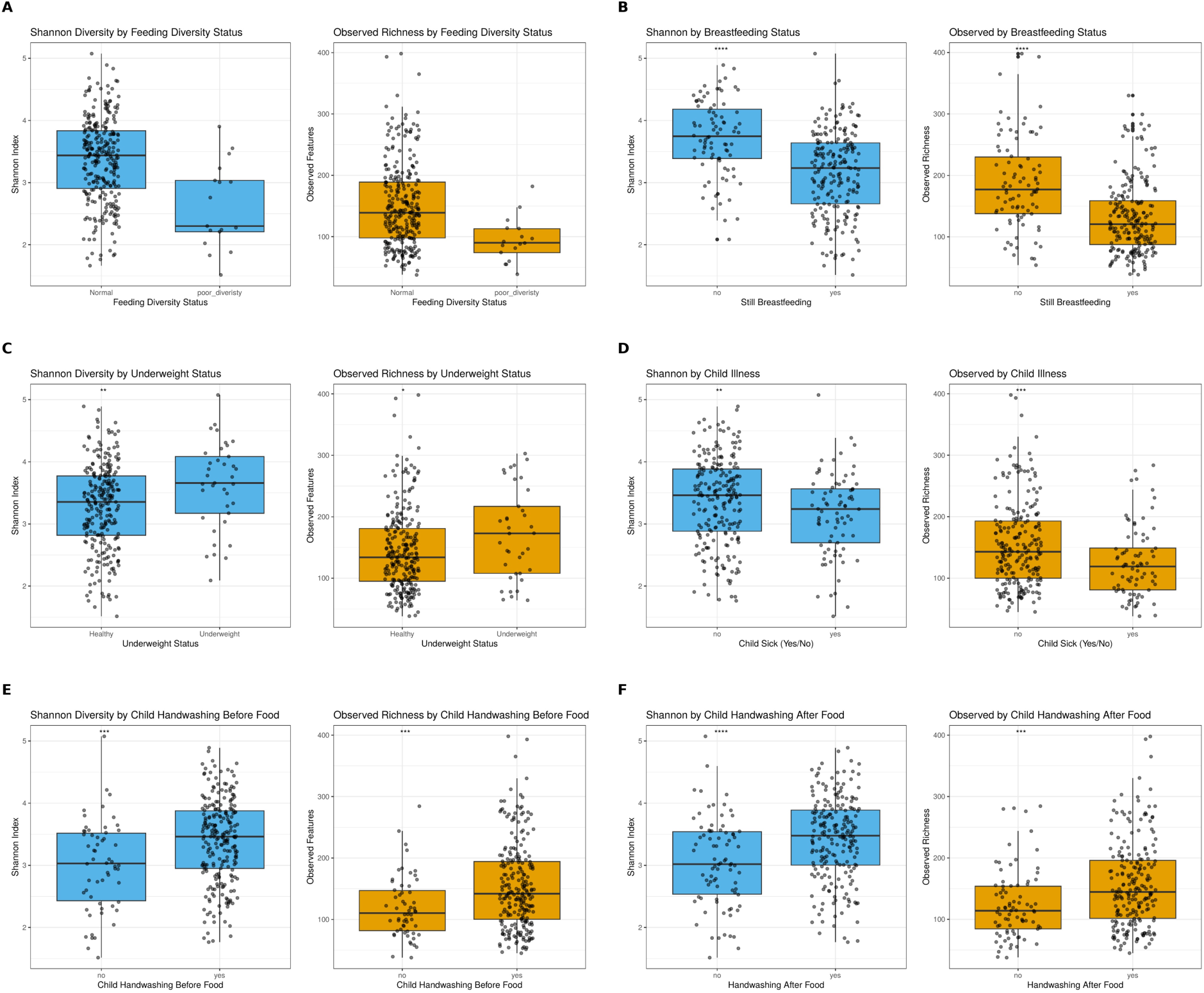
Association between covariates and gut microbial diversity. Boxplots display variations in alpha diversity indices. Statistical differences in alpha diversity were tested using Kruskal Wallis test, with p-values < 0.05 considered significant. (A) dietary diversity status (B) Extended breastfeeding (C) Underweight status (D) child illness (E) child handwashing before a meal (F) child handwashing after a meal.

Furthermore, although no clear separation can be observed, PERMANOVA shows significantly different microbial community composition with dietary diversity status (p = 0.001), underweight status (p=0.005), children whose hands were washed by their caregivers before or after feeding (p = 0.001), child illness (p = 0.022), and extended breastfeeding (p = 0.001) (figure 5).

**Figure 5:**
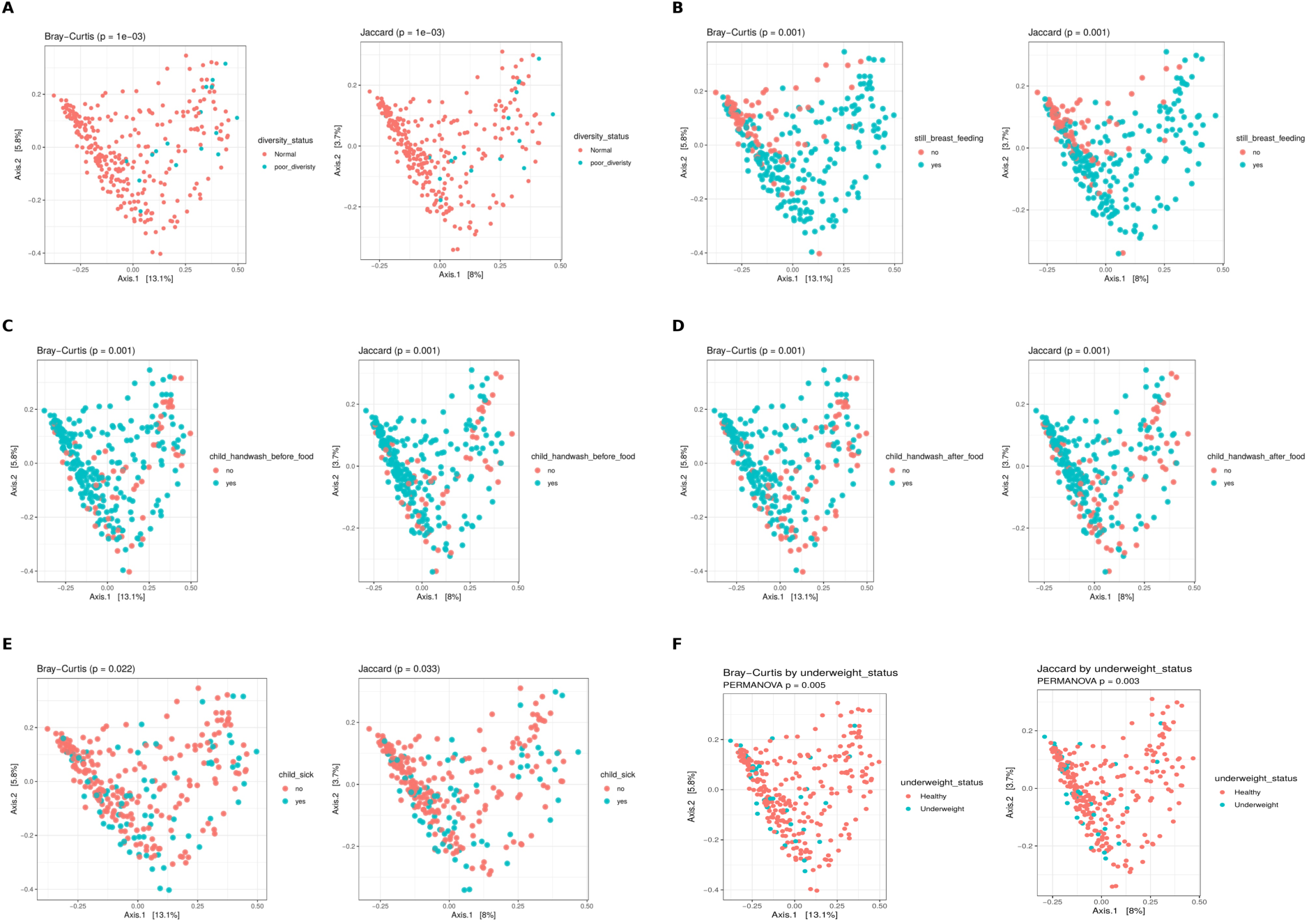
Gut microbiome composition by significant explanatory variables. From principal coordinates analysis (PCoA) plots, points represent individual samples colored by group status. Group differences in community composition were tested using PERMANOVA, with p-values indicated in the ordination plots. Axis labels show the percentage of variance explained. (A) dietary diversity status (B) Extended breastfeeding (C) Underweight status (D) child illness (E) child handwashing before a meal (F) child handwashing after a meal.

The multi-site centers selection revealed location variations in microbiota composition and diversity. Geographical residence also reported significant differences in microbial diversity across districts (p = 0.044) and wards (p = 0.0007) as well as microbial variation across districts (p =0.008) in supplementary figure 2. For instance, children in Kilolo district were associated with greater microbial richness (p = 0.0045).

In this study, gender, gestation period, child delivery method, a recent travel outside the residence place, toilet sharing among households within the same structure, and caregiver hand washing practices before and after feeding the child were not a major driver of microbial diversity and composition difference (p > 0.05).

### 3.4 Taxonomic composition and differential abundance

We identified a total of 20 bacterial phyla and 437 genera from fecal samples of children. Among stunted children, 395 bacterial genera were detected, while for children with normal growth the number of genera detected was 283. From the 20 phyla detected (Table 2), 18 were shared by both groups. Among the detected genera, the top 30 most abundant taxa dominated the gut bacteriome collectively. *Prevotella* was the most dominant genus (mean relative abundance 21.7%), followed by *Bifidobacterium* (12.2%) and *Faecalibacterium* (10.8%) (Figure 6).

**Figure 6:**
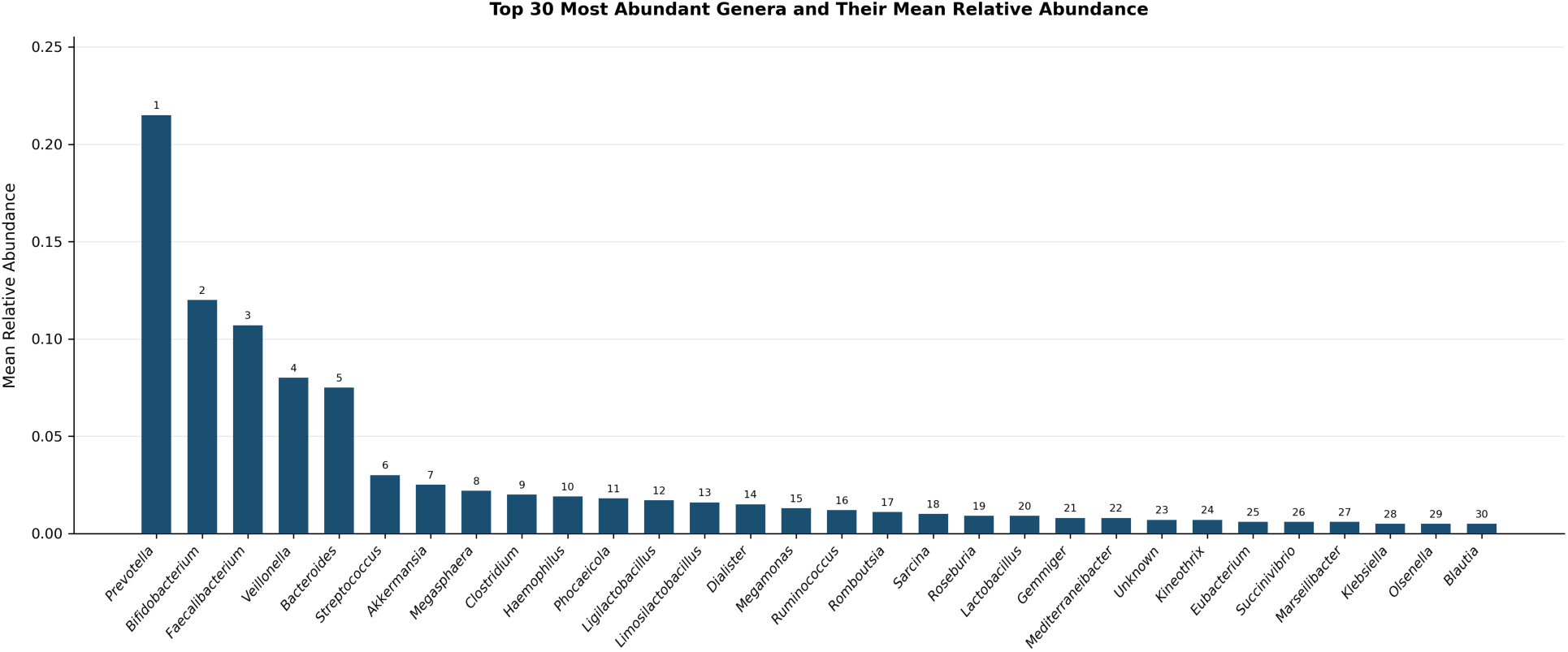
Top 30 most abundant bacterial genera in the study population. Bar chart displaying the mean relative abundance of the 30 most prevalent gut bacterial genera identified across all samples. Genera are ranked in descending order of mean relative abundance (rank indicated above each bar).

**Table 2:** Total bacterial phyla detected across all samples and by stunting status in the study population.

| Sn | Phylum | Number of taxa | Percentage (%) | Detected in stunted | Detected in normal |
| --- | --- | --- | --- | --- | --- |
| 1 | Firmicutes | 7812 | 53.29 | ✓ | ✓ |
| 2 | Bacteroidetes | 3441 | 23.47 | ✓ | ✓ |
| 3 | Proteobacteria | 2133 | 14.55 | ✓ | ✓ |
| 4 | Actinobacteria | 868 | 5.92 | ✓ | ✓ |
| 5 | Verrucomicrobia | 146 | 1.00 | ✓ | ✓ |
| 6 | Fusobacteria | 115 | 0.78 | ✓ | ✓ |
| 7 | Unknown | 36 | 0.25 | ✓ | ✓ |
| 8 | Cyanobacteria | 33 | 0.23 | ✓ | ✓ |
| 9 | Bacteroidetes-Firmicutes | 18 | 0.12 | ✓ | ✓ |
| 10 | Tenericutes | 18 | 0.12 | ✓ | ✓ |
| 11 | Acidobacteria | 8 | 0.05 | ✓ | ✓ |
| 12 | Spirochaetes | 7 | 0.05 | ✓ | ✓ |
| 13 | Planctomycetes | 5 | 0.03 | ✓ | ✓ |
| 14 | Gemmatimonadetes | 4 | 0.03 | ✓ | ✓ |
| 15 | Synergistetes | 4 | 0.03 | ✓ | ✓ |
| 16 | Chlamydiae | 3 | 0.02 | ✓ | ✓ |
| 17 | Chloroflexi | 3 | 0.02 | ✓ | ✗ |
| 18 | Elusimicrobia | 3 | 0.02 | ✓ | ✓ |
| 19 | Candidatus_Saccharibacteria | 2 | 0.01 | ✓ | ✓ |
| 20 | Chlamydiae-Firmicutes | 1 | 0.01 | ✗ | ✓ |

Table 2 shows distribution of bacterial phyla identified from 16S rRNA gene sequencing across the study cohort. The table summarizes all bacterial phyla detected across all samples, as well as phyla detected among children with stunted growth and children with normal growth. Phyla were identified following taxonomic classification of quality-filtered sequences.

A total of 25 core bacterial genera were identified across all children, defined as genera present in at least 50% of samples with a mean relative abundance ≥ 0.1%, with 24 genera shared with stunted children and only 16 with normally growing children. *Megasphaera* was absent among children with stunted growth while *Ligilactobacillus, Romboutsia, Intestinibacter, Roseburia, Dorea, Kineothrix, Parabacteroides, Ruminococcus* and *Eubacterium* were absent in children with normal growth (Figure 7).

**Figure 7:**
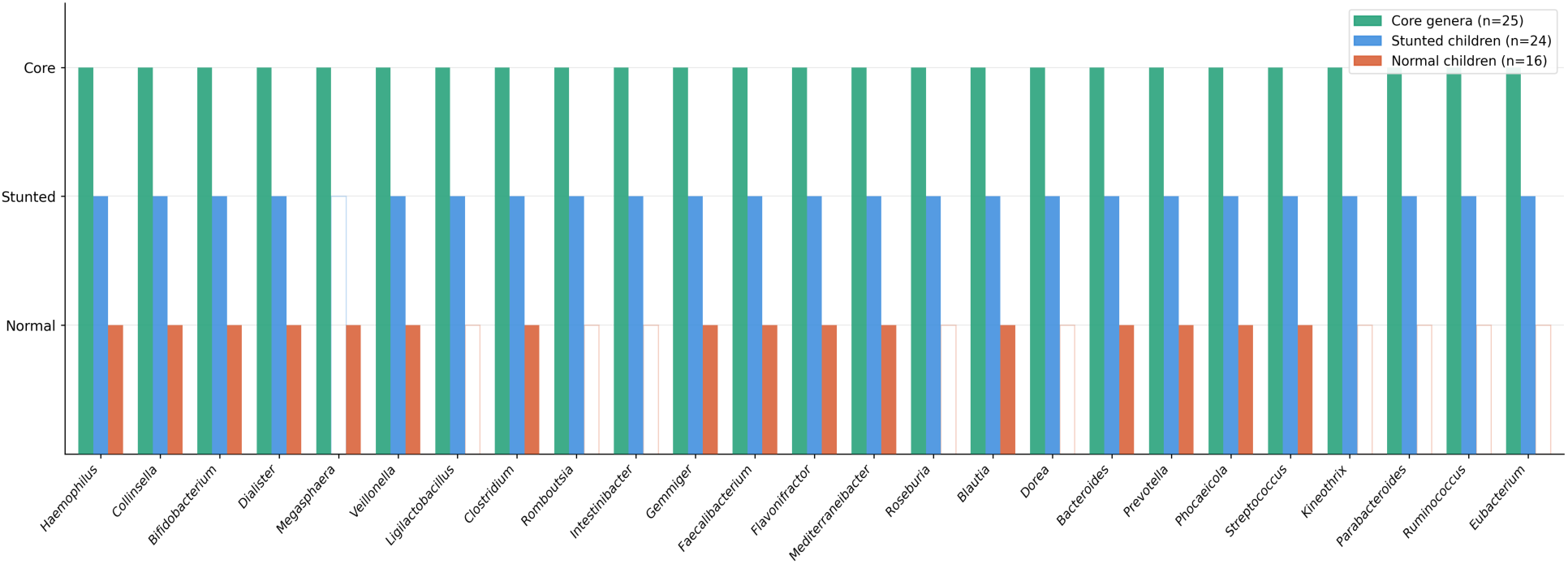
Core bacterial genera detected across all fecal samples of the sampled children. Blue bars indicate the core genera in stunted children and red bars in normal growing children.

#### Taxonomic composition for covariates

Table 3 summarizes the taxonomic classification of the identified taxa and number of bacterial phyla and genera detected in fecal samples according to dietary practices, underweight status, extended breastfeeding, recent child illness, hygiene behaviors through handwashing practices, and ward of residence. A taxon was considered detected if it was present in at least one sample within the respective subgroup. Counts therefore reflect presence within a category and do not indicate relative abundance or statistical significance.

**Table 4:**
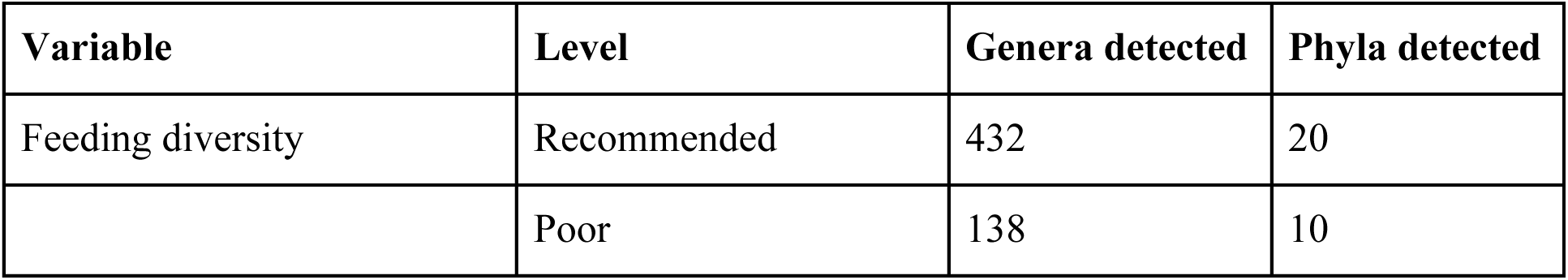
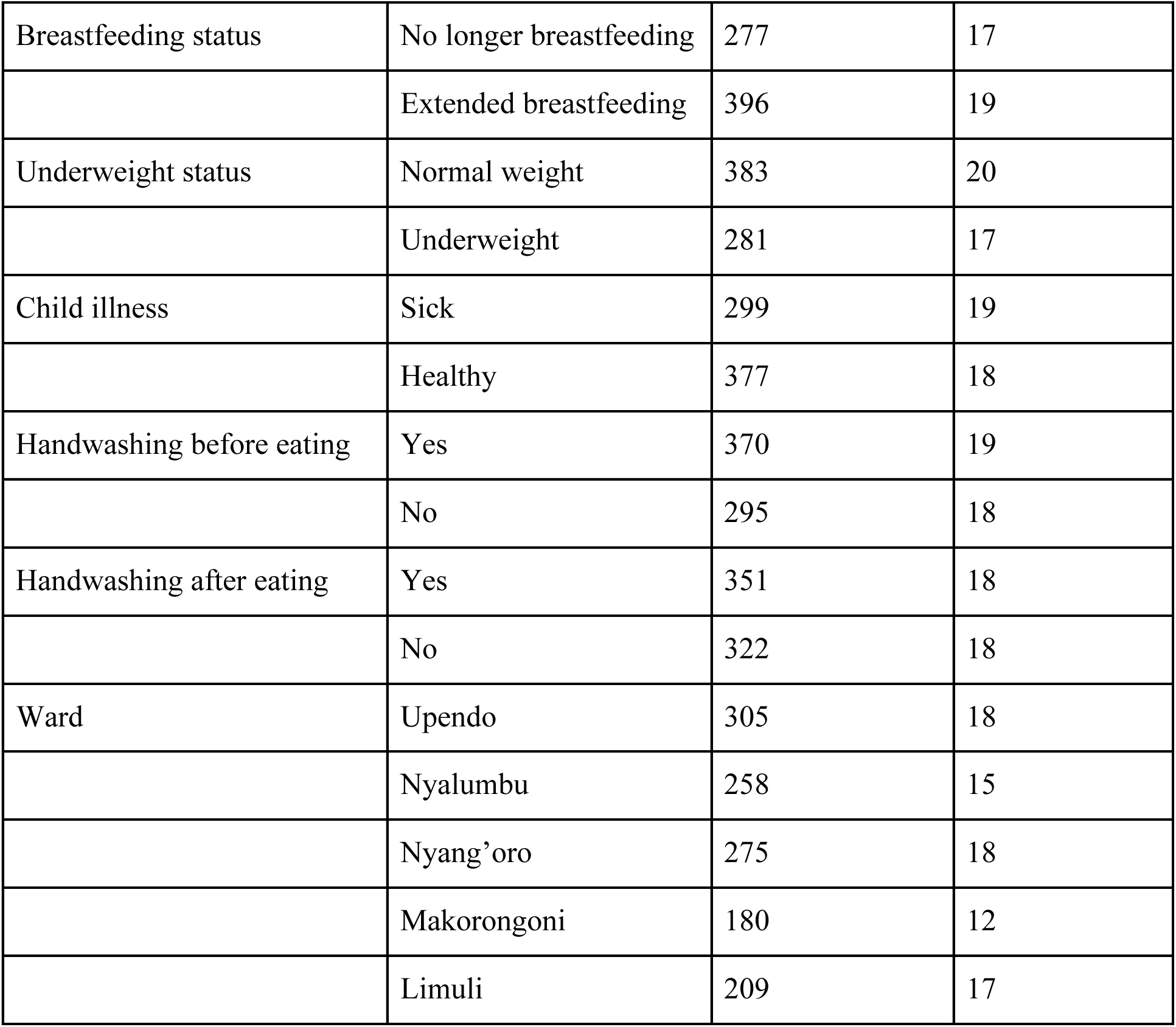
A summary of detected taxa across exposure variables.

| Variable | Level | Genera detected | Phyla detected |
| --- | --- | --- | --- |
| Feeding diversity | Recommended | 432 | 20 |
|  | Poor | 138 | 10 |
| Breastfeeding status | No longer breastfeeding | 277 | 17 |
|  | Extended breastfeeding | 396 | 19 |
| Underweight status | Normal weight | 383 | 20 |
|  | Underweight | 281 | 17 |
| Child illness | Sick | 299 | 19 |
|  | Healthy | 377 | 18 |
| Handwashing before eating | Yes | 370 | 19 |
|  | No | 295 | 18 |
| Handwashing after eating | Yes | 351 | 18 |
|  | No | 322 | 18 |
| Ward | Upendo | 305 | 18 |
|  | Nyalumbu | 258 | 15 |
|  | Nyang'oro | 275 | 18 |
|  | Makorongoni | 180 | 12 |
|  | Limuli | 209 | 17 |

#### Bacterial differential abundance

Children with normal growth were enriched in *Bifidobacterium, Rothia, Olsenella, Slackia, Lactobacillus, Gemella,* and *Oscillibacter,* while these taxa were suppressed in stunted children. On the other side, stunted children were enriched in the *Ruminococcus torques group, Acinetobacter, Prevotella, Alistipes, Odoribacter, Fournierella, Fusobacterium,* and *Akkermansia* (Figure 8).

**Figure 8:**
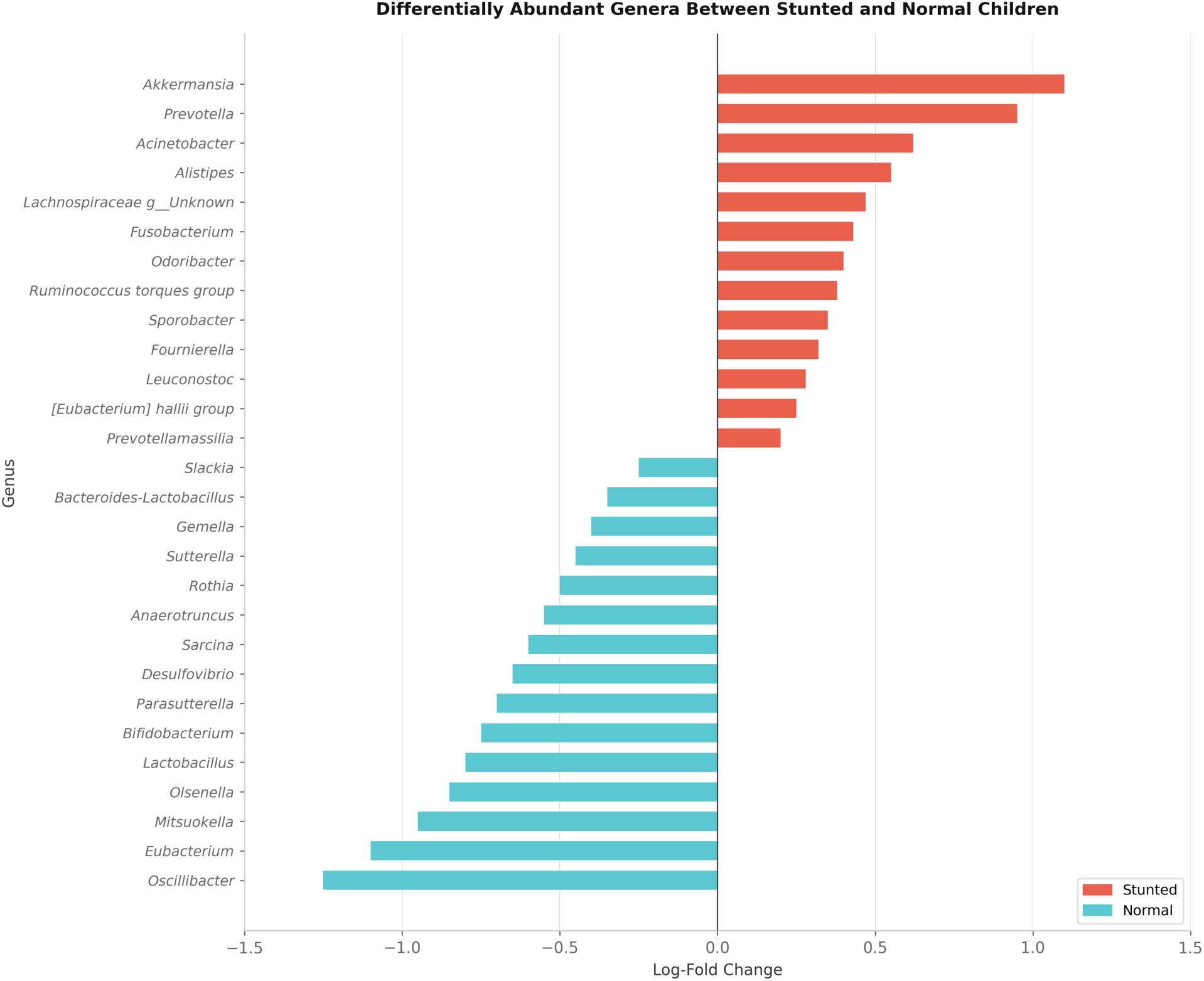
Differentially abundant bacterial genera for stunting status showing significant differences in relative abundance between stunted and normal growing children. Bars represent log-fold changes in genus abundance, with red bars indicating enrichment in the stunted group and blue bars indicating enrichment in the normal group

Among the 40 tested genera for dietary diversity status, 5 genera were enriched in children with poor feeding diversity status measured through minimum dietary diversity score, while 4 genera were marked in children with better feeding diversity (Figure 9A). Children with recent illness showed enrichment of opportunistic and gut genera potential for causing inflammation while healthy children were enriched in beneficial taxa (Figure 9B). A total of 9 taxa characterized by genera linked to fermentation dynamics and potential environmental exposure were enriched among children from households sharing toilets, whereas 14 genera exhibiting a more diverse set of taxa were enriched among children from single-structure households (Figure 9C).

**Figure 9:**
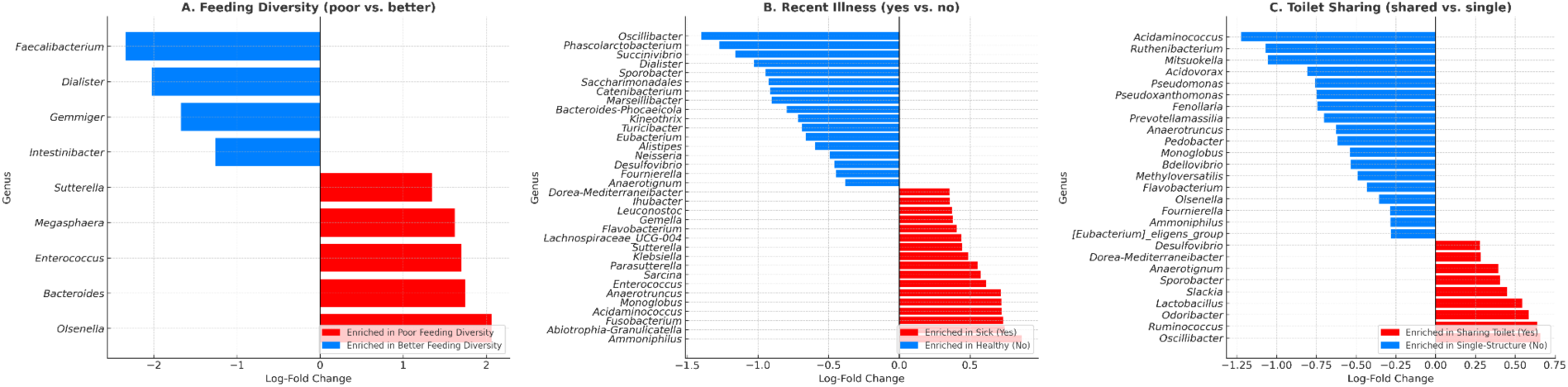
Differentially abundant bacterial genera across household factors. children. (A) Differentially abundant genera associated with feeding diversity (poor vs. better). (B) Genera differing by recent illness status (yes vs. no). (C) Genera differing by toilet sharing (shared vs. single). Bars represent log-fold changes in genus abundance, with red bars indicating enrichment in the first group of each comparison (poor feeding diversity, recent illness, shared toilet) and blue bars indicating enrichment in the contrasting group.

### 3.5 Predictive Modeling of Stunting

A subset of microbial genera was identified as significant predictors of stunting status with 87.5% accuracy based on 10-fold cross-validation. Key genera identified as predictors of stunted growth were *Veillonella* (importance = 0.039), *Rothia* (importance = 0.028), *Streptococcus* (importance = 0.032), and *Clostridium* (importance = 0.030), while *Bifidobacterium* (importance = 0.045) and *Faecalibacterium* (importance = 0.035) showed stronger associations with normal growth. An unclassified taxon (Unclassified, importance = 0.25) also ranked highly as a predictor (supplementary figure 2).

## 4.0 Discussion

### 4.1 Demographics of the study population

This study found that six out of ten children are stunted in Iringa. Significant associations between stunting and child age, gender, continued breastfeeding, dietary diversity, household toilet sharing, geographic residence, and child handwashing practices were identified. Lefebo and colleagues reported a similar association between stunting and breastfeeding duration as well as residence (32). Stunted children were older (mean 15.7 months) than normal growing children (mean 13.1 months) (p < 0.001), suggesting prolonged exposure to risk factors, consistent with other studies (33). The age association with stunting is comparable to results from Batool and colleagues (6).

Stunted children were more likely to be male (57%). The higher prevalence of stunting in boys (p = 0.049) aligns with other findings of gender-specific vulnerabilities (34,35). Some studies report higher stunting rates in boys are potentially influenced by their higher energy needs (36).

Continued breastfeeding was protective of stunting (p=0.001), reinforcing WHO recommendations for breastfeeding up to 2 years. Households who shared toilet facilities (p =0.001) and child handwashing before and after eating (p =0.003, & p =0.007) were also associated with stunting status, possibly reflecting improved sanitation practices or health interventions in the community. Child delivery mode and recent child illness showed no significant association with stunting (p > 0.05), showing that these factors may not play a major role in this population. Similar findings were reported, where more stunted children were born by vaginal delivery compared to caesarian, however the logistic model demonstrated no significant impact of mode of delivery on stunting status (37).

### 4.2 Gut microbial diversity and stunting

We identified differences in the fecal bacterial diversity and composition between normally growing and stunted children. Stunted children exhibited higher alpha diversity, a pattern observed to be primarily attributable to age-related gut microbiome maturation rather than a direct effect of stunting. On average, stunted children were older (mean 15.7 months) than those with normal growth (mean 13.1 months), with a strong association between age and microbial diversity and composition driving this trend. This aligns with prior research reporting microbial community stabilization during early childhood (38). In contrast, a longitudinal study reported lower gut microbiota diversity in stunted versus healthy children, though it similarly highlighted developmental changes influenced by age (39).

Statistically, the gut bacterial community structure of stunted children differed significantly from that of normal growing children, as evidenced by PERMANOVA, similar to other results (40). The community structure interspersed with no clear distinct clustering, meaning that bacterial community composition is largely similar with no strong compositional shift between stunted and normal growing children based on presence or absence of a taxa. These differences were strongly influenced by age, dietary practices, child sickness and environmental factors, including WaSH. Despite the region’s food-secure status where stunting remains high, this suggests the role of combined (nutritional and environmental) drivers. Consistent with these findings, a recent study from East Nusa Tenggara, Indonesia, using 16S rRNA sequencing, reported significant compositional differences between stunted and normal growing children, attributing these to environmental factors like sanitation and dietary intake (41).

### 4.3 Taxonomic composition and abundance

Our study population displayed 20 bacterial phyla and 437 genera across all fecal samples. Notably, stunted children harbored more (395) genera compared to in normal growing children (283 genera), while most phyla were shared in both groups. This substantial increase in genus-level diversity among stunted children may suggests that stunting is associated with a broader taxonomic range, potentially further suggesting colonization by additional opportunistic or environmentally derived taxa.

Among the top 30 most abundant genera, *Prevotella* was the most dominant taxon, followed by *Bifidobacterium* and *Faecalibacterium*. The dominance of *Prevotella* is characteristic of plant-rich and high-fiber diets and has been associated with fibre fermentation and SCFA production. However, certain *Prevotella* species have also been implicated in intestinal inflammation (42). Other taxa included *Blautia, Dialister,* and *Lactobacillus*, indicating a complex and evolving microbial structure during this critical developmental window (first 1000 days). This diversity suggests a baseline resilient structure, yet susceptibility to environmental and nutritional exposures as similarly reported by Dinh et al (43).

Also, our study revealed a diverse yet shared core gut bacteriome among the study cohort. Several of the core genera are well-established short-chain fatty acids (SCFAs) producers mostly butyrate and propionate. *Faecalibacterium*, *Roseburia*, *Blautia*, *Ruminococcus*, and *Eubacterium* are among the most important butyrate producers in the human gut (44). Butyrate is the primary energy source for colonocytes and plays a critical role in maintaining gut barrier integrity and suppressing intestinal inflammation. *Prevotella* and *Dialister* are known propionate producers through succinate pathways (45). The presence of these genera in both groups suggests a baseline capacity for SCFA production, though the relative abundance differences between stunted and normal children would determine the magnitude of this functional output.

*Clostridium* and *Intestinibacter* are capable of mucin degradation, which at normal levels supports mucus layer turnover but when elevated may compromise the mucus barrier (46)(47). *Bifidobacterium*, *Faecalibacterium*, and *Blautia* are strongly associated with anti-inflammatory activity. *Bifidobacterium* promotes regulatory T-cell development and produces acetate which reinforces epithelial barrier function. *Faecalibacterium prausnitzii* is one of the most studied anti-inflammatory gut bacteria, suppressing NF-κB signalling and IL-8 production (48). Their presence in the core of both groups is reassuring.

*Clostridium*, *Streptococcus*, and *Veillonella* have species with either role, commensal at low abundance but potentially pro-inflammatory when dominant. *Veillonella* metabolises lactate from primary fermenters and produces propionate, but some species have been associated with systemic inflammation. *Streptococcus* in the gut has been linked to intestinal permeability when in excess (49). *Collinsella*, *Ligilactobacillus*, and *Bifidobacterium* produce lactate and acetate, contributing to a lowered gut pH that inhibits pathogen growth (50). This is particularly important in young children. Presence of these most prominent bacterial genera as supported by literature (51), suggests their benefits to the host.

Children with normal growth were enriched in beneficial genera, including *Bifidobacterium, Rothia, Olsenella, Slackia, Lactobacillus, Gemella,* and *Oscillibacter,* aligning with findings from Gatya et al. (2022) (52), who reported depletion of these taxa in stunted children. In contrast, stunted children showed higher abundance of *Ruminococcus torques group, Acinetobacter, Prevotella, Alistipes, Odoribacter, Fournierella, Fusobacterium,* and *Akkermansia* as also reported by (53). As it has been reported, stunted children often face prolonged exposure to environmental stressors like poor WaSH, recurrent infections including diarrhea, or suboptimal feeding (54). These factors may introduce additional low-abundance or opportunistic taxa; inflating genus counts without altering core phyla.

The differential enrichment of SCFA producers (*Bifidobacterium*, *Faecalibacterium*, *Oscillibacter*, *Eubacterium*, *Anaerotruncus*) in normal-growth children suggests a greater capacity for SCFA production including butyrate, a key factor in promoting nutrients absorption for growth (55,56). Conversely, the high abundance of *Akkermansia*, *Ruminococcus* torques group, mucin degraders, alongside *Prevotella* in stunted children, may indicate a disrupted mucus layer, potentially exacerbating inflammation, though this requires species level and functional validation (57–59). A recent study from Pemba, Tanzania, using 16S rRNA sequencing, similarly identified *Akkermansia*, *Prevotella*, and *Odoribacter* as strong markers of nutritional status, though the direction of association varied by context (60). These findings underscore the complex interplay of microbial composition and environmental exposures in shaping stunting outcomes.

When considering the weight for age of the children, the gut composition showed a higher richness among healthy children (383 genera) compared with those underweight (281 genera). Undernutrition channeled from insufficient nutrients required for growth may reduce microbial diversity through lower dietary quantity and quality, impaired gut barrier function, and repeated infections (61).

### 4.4 Dietary influences on Microbiota

Children with recommended dietary diversity exhibited higher alpha diversity, including richness, possibly reflecting the consumption of a variety of food sources that support diverse microbial niches. Similar findings were reported by Laitinen and Mokkala (62), where pregnant women with highest dietary quality had higher gut microbiota diversity in all the investigated indexes. On the other hand, poor dietary diversity was associated with altered bacterial community structures, probably because diverse diet choices fostered a distinct microbial profile compared to monotonous diets as reported by parents and guardians. This aligns with findings from another study, where higher diet quality was linked to increased alpha diversity and beta composition (63).

In this study, children with recommended dietary diversity had broader bacteriome characterized in most (432) genera compared to less (138) genera distributed across less phyla found in children with poor dietary diversity. The markedly lower number of detected taxa in the poor feeding diversity group suggests a reduced breadth of microbial representation compared with children consuming more diverse diets. Literature suggests that monotonous diets may restrict the range of ecological niches. Diverse diets typically provide a mixture of fibers that support multiple microbial niches and promote a more stable community (64).

Children with better dietary diversity were enriched in *Faecalibacterium, Gemmiger,* and *Dialister* genera, known for their robust butyrate production. On the other hand, those with poor feeding diversity showed enrichment in *Olsenella, Enterococcus*, and *Megasphaera*, suggesting that limited dietary variety may promote fewer desirable taxa. Our findings aligns with those of De Filippo et al. (2010), where a high-fiber diet enriched *Prevotella* and *Xylanibacter* fiber degraders linked to increased SCFAs while a western diet was associated with Enterobacteriaceae, supporting the role of dietary diversity in shaping microbial composition and health outcomes (65).

### 4.5 Extended breastfeeding and Microbial diversity

Extended breastfeeding beyond six months of exclusive breastfeeding was associated with lower alpha diversity in this cohort, consistent with literature evidence that early-life microbial communities, dominated by *Bifidobacterium* due to breastmilk. Transiting to higher diversity follows complementary food introduction and/or breastfeeding cessation (66). Nevertheless, children who were still breastfeeding harbored more (396) genera, compared with fewer (277) genera in those no longer breastfed. Additionally, extended breastfeeding was associated with enrichment in protective genera such as *Bifidobacterium* and *Lactobacillus*, which may mitigate the effects environmental pathogenic exposures due to early complementary feeding and by fostering early gut colonization as reported by Differding and colleagues (67). However, children who were no longer breastfeeding exhibited higher levels of *Prevotella* and *Roseburia*, alongside other beneficial taxa, indicating a rapid shift toward a mature microbial profile.

Continued exposure to mother’s milk likely supplies the child with human milk oligosaccharides (HMO) that nourish beneficial microbes (68). Also, breast milk provides antibodies that fight against pathogens and inflammation, which helps maintain ecological space for a richer microbiota (69). This shift underscores the need to balance breastfeeding duration with diverse feeding practices to optimize microbial diversity in early childhood (70).

### 4.6 Environmental factors and geographical residence

Geographical residence influenced microbial profile differences, with children in Kilolo district showing elevated microbial richness, possibly due to unique dietary practices, warranting further investigation into environmental drivers, similar to Senghor (71). This highlights the need for targeted interventions in high-risk areas.

Regular handwashing by parents/guardians and children before and after meals was associated with higher diversity, suggesting reduced pathogen exposure and improved hygiene practices, a finding supported by Ratnayani et al. (2024) (72), who linked hygiene and macronutrient intake to microbial diversity. However, our observations contrast with the SHINE trial, where WaSH and feeding interventions did not significantly alter fecal microbiota or influence growth via the microbiome (73).

Moreover, more (370) were genera detected in children who were hand washed by their caregivers before feeding compared to where this practice was absent (295). A similar, though smaller, pattern appeared for handwashing after eating (351 vs. 322 genera). Hygiene likely reduces repeated exposure to enteropathogen and contamination, thereby protecting commensal organisms (74).

Conversely, acute illness at the time of the survey correlated with lower diversity across indices and altered composition, suggesting microbial disruption. This is consistent with other findings, where diarrheal frequency and severity were negatively associated with bacterial diversity and richness, with stunted children exhibiting greater diversity losses and slower recovery (75).

The observed abundance from sick children was lower (299 genera) compared to healthy children suggests that illness, diarrhea, can cause dysbiosis leading to loss of sensitive commensals and overgrowth of opportunists microbes. Reductions of these taxa suggest loss of reasonable diversity (76).

Opportunistic genera including *Enterococcus*, *Fusobacterium*, and *Klebsiella*, were enriched in children reported to have been recently ill, suggesting that infection may induce a state of dysbiosis that could impair nutrient absorption, a known risk factor for stunting. This aligns with Reyman et al. (2022) (77), who observed increased *Klebsiella* and *Enterococcus spp*., alongside reduced *Bifidobacterium spp.*, in antibiotic-treated neonates, indicating a shared microbial response to infection or treatment that may exacerbate growth deficits.

### 4.7 Predictive taxa of Stunting

We have found genera linked to a healthy gut, including *Bifidobacterium* and *Faecalibacterium*, to be strong predictors of normal growth, reflecting their roles in gut maturation and SCFA production, carbohydrate fermentation, and immune modulation during early childhood while *Pretovella*, *Veillonella*, *Rothia*, *Streptococcus*, and *Clostridium* were predictive of stunting, suggesting maybe a dysbiotic state potentially tied to inflammation or pathogen exposure. These taxa, validated by their high feature importance, serve as robust microbial markers for stunting status. The model’s identification of an unclassified taxon as a significant predictor underscores the potential contribution of novel or poorly characterized species, likely undetected by targeted 16S rRNA sequencing. This finding supports the development of a microbiome-based diagnostic tool, offering a non-invasive screening approach to identify at-risk children, though its clinical utility requires validation through larger studies. Future research employing whole-genome sequencing is essential to characterize these unclassified taxa and refine predictive accuracy for stunting interventions.

### 4.8 Study limitations

This study has several limitations that should be considered. The cross-sectional design limits the ability to establish causality between gut bacteriome profiles and stunting, as it captures a single time point rather than longitudinal changes. Additionally, gut microbiome characterization was performed using targeted amplicon sequencing of the V3–V4 hypervariable regions of the 16S rRNA gene. While this approach is well-validated, widely used, and appropriate for community-level profiling, it has inherent resolution limits for reliable species-level taxonomic assignments, as multiple bacterial species can share near-identical 16S rRNA sequences. Accordingly, all taxonomic assignments and biological interpretations in this study are reported and discussed at the genus level. Species-level characterisation and functional metabolic pathway analysis will be addressed in subsequent shotgun metagenomic sequencing of the same cohort, which is currently underway. Potential recall bias in household surveys, particularly for dietary and hygiene practices, may affect the accuracy of metadata. Nutritional exposures were only assessed through, dietary quantity and micronutrient intake were not assessed, limiting full interpretation of nutrition-related determinants

Unmeasured confounders, such as maternal health, could also influence results. The community-based sampling may introduce selection bias, potentially limiting generalizability to other regions or populations, although the use of randomization software was used to minimize the risk. Future longitudinal studies with objective measures and whole genome analyses are recommended to address these limitations and confirm the findings.

## 5.0 Conclusion

This study aimed to explore the gut bacteriome profiles of children in Iringa, Tanzania, as a potential biomarker for growth and to assess the influence of environmental factors, particularly water, sanitation, and hygiene (WaSH) and feeding practices. Through a targeted V3-V4 of the 16S rRNA sequencing complemented by household surveys, findings revealed that stunting, affecting 60.5% of children, was associated with microbial profiles. A shared core bacteriome is observed in both groups, hence stunting does not appear to be caused by a completely different gut microbiome. Instead, shared environmental and dietary factors shape both the gut bacteria and child growth. Normal growing children were enriched in beneficial genera like *Bifidobacterium, Lactobacillus* and *Faecalibacterium*, while stunted children showed higher *Prevotella* and *Akkermansia*. Age was the most consistent predictor of gut microbial diversity whereas dietary diversity, continued breastfeeding, hygiene practices, recent illness and residential location also significantly influenced microbial diversity and composition. These results underscore the need for interventions promoting diverse complementary feeding along with continued breastfeeding and improved WaSH practices during the critical growth window to reduce stunting in Iringa and similar regions.

## Authors contribution

GM and KSM participated in brainstorming the idea. GM initiated writing the first draft of the manuscript. All authors (AD, BL, SL, IA and KSM) participated in writing: reviewing and editing, specifically critical review, commentary or revision. All authors have approved the manuscript for submission.

## Acknowledgement

The authors would like to sincerely thank the parents/guardians and community members in Iringa region who participated in this study. We acknowledge the support of local leaders and community health workers for their assistance in sensitization and mobilizing households during data collection. We are also grateful to the research assistants and field team members for their dedication throughout the survey. Special appreciation goes to the Nelson Mandela African Institution of Science and Technology (NM-AIST) and the University of Dar es Salaam (UDSM) for providing institutional support. This study was funded by the Nestlé Foundation for the study of problems of nutrition in the world and the 100 PhDs for Africa programme under the UM6P – EPFL Excellence in Africa Initiative, whose support made this research possible.

## References

1. World Health Organization, United Nations Children’s Fund. Global nutrition targets 2030: stunting brief. World Health Organization; 2025.

2. World Health Organization. Reducing stunting in children: Equity considerations for achieving the Global Nutrition Targets 2025. Geneva; 2018.

3. UNICEF, W H O, The World Bank. Joint Child Malnutrition Estimates (JME): Levels and trends – 2025 edition. Geneva: World Health Organization; 2025 Jul.

4. Elisaria E, Caeyers B, Nkuba E, van der Erve L, Kuwawenaruwa A. Thirty years of declining stunting in Tanzania: Trends and ongoing challenges. PLoS ONE. 2025 Jul 29;20(7):e0327779.

5. Makundi, A, Ndekia TE. Road to accelerated reduction of childhood stunting in Tanzania: The Njombe story [Internet]. UNICEF Tanzania. 2024 [cited 2025 Oct 27]. Available from: https://www.unicef.org/tanzania/stories/road-accelerated-reduction-childhood-stunting-tanzania

6. Batool M, Saleem J, Zakar R, Butt MS, Iqbal S, Haider S, et al. Relationship of stunting with water, sanitation, and hygiene (WASH) practices among children under the age of five: a cross-sectional study in Southern Punjab, Pakistan. BMC Public Health. 2023 Nov 3;23(1):2153.

7. Ministry of Health (MoH) [Tanzania Mainland], Ministry of Health (MoH) [Zanzibar], National Bureau of Statistics (NBS), Office of the Chief Government Statistician (OCGS), ICF. Tanzania Demographic and Health Survey and Malaria Indicator Survey 2022: Key Indicators Report. Dodoma, Tanzania, and Rockville, Maryland, USA: MoH, NBS, OCGS, and ICF; 2023.

8. Pickard JM, Zeng MY, Caruso R, Núñez G. Gut microbiota: Role in pathogen colonization, immune responses, and inflammatory disease. Immunol Rev. 2017 Sep;279(1):70–89.

9. Kamada N, Chen GY, Inohara N, Núñez G. Control of pathogens and pathobionts by the gut microbiota. Nat Immunol. 2013 Jul;14(7):685–90.

10. Rodríguez JM, Murphy K, Stanton C, Ross RP, Kober OI, Juge N, et al. The composition of the gut microbiota throughout life, with an emphasis on early life. Microb Ecol Health Dis. 2015 Feb 2;26:26050.

11. Nunez H, Nieto PA, Mars RA, Ghavami M, Sew Hoy C, Sukhum K. Early life gut microbiome and its impact on childhood health and chronic conditions. Gut Microbes. 2025 Dec;17(1):2463567.

12. Wang H, Wei C-X, Min L, Zhu L-Y. Good or bad: gut bacteria in human health and diseases. Biotechnology & Biotechnological Equipment. 2018 Jun 11;1–6.

13. Cheng J, Ringel-Kulka T, Heikamp-de Jong I, Ringel Y, Carroll I, de Vos WM, et al. Discordant temporal development of bacterial phyla and the emergence of core in the fecal microbiota of young children. ISME J. 2016 Apr;10(4):1002–14.

14. Fobofou SA, Savidge T. Microbial metabolites: cause or consequence in gastrointestinal disease? Am J Physiol Gastrointest Liver Physiol. 2022 Jun 1;322(6):G535–52.

15. Syromyatnikov M, Nesterova E, Gladkikh M, Smirnova Y, Gryaznova M, Popov V. Characteristics of the gut bacterial composition in people of different nationalities and religions. Microorganisms. 2022 Sep 18;10(9).

16. Chac D, Slater DM, Guillaume Y, Dunmire CN, Ternier R, Vissières K, et al. Association between chlorine-treated drinking water, the gut microbiome, and enteric pathogen burden in young children in Haiti: An observational study. Int J Infect Dis. 2024 Oct;147:107165.

17. Monira S, Barman I, Jubyda FT, Ali SI, Islam A, Rahman KMZ, et al. Gut microbiota shifts favorably with delivery of handwashing with soap and water treatment intervention in a prospective cohort (CHoBI7 trial). J Health Popul Nutr. 2023 Dec 21;42(1):146.

18. Regassa R, Tamiru D, Duguma M, Belachew T. Environmental enteropathy and its association with water sanitation and hygiene in slum areas of Jimma Town Ethiopia. PLoS ONE. 2023 Jun 23;18(6):e0286866.

19. Dagar S, Singh J, Saini A, Kumar Y, Chhabra S, Minz RW, et al. Gut bacteriome, mycobiome and virome alterations in rheumatoid arthritis. Front Endocrinol (Lausanne). 2022;13:1044673.

20. Peroni DG, Nuzzi G, Trambusti I, Di Cicco ME, Comberiati P. Microbiome composition and its impact on the development of allergic diseases. Front Immunol. 2020 Apr 23;11:700.

21. Bintsis T. Foodborne pathogens. AIMS Microbiol. 2017 Jun 29;3(3):529–63.

22. Sanin KI, Haque A, Nahar B, Mahfuz M, Khanam M, Ahmed T. Food Safety Practices and Stunting among School-Age Children-An Observational Study Finding from an Urban Slum of Bangladesh. Int J Environ Res Public Health. 2022 Jun 30;19(13).

23. Budge S, Parker AH, Hutchings PT, Garbutt C. Environmental enteric dysfunction and child stunting. Nutr Rev. 2019 Apr 1;77(4):240–53.

24. Xu P, Lv T, Dong S, Cui Z, Luo X, Jia B, et al. Association between intestinal microbiome and inflammatory bowel disease: Insights from bibliometric analysis. Comput Struct Biotechnol J. 2022 Apr 7;20:1716–25.

25. Martín R, Rios-Covian D, Huillet E, Auger S, Khazaal S, Bermúdez-Humarán LG, et al. Faecalibacterium: a bacterial genus with promising human health applications. FEMS Microbiol Rev. 2023 Jul 5;47(4).

26. Ghosh TS, Shanahan F, O’Toole PW. The gut microbiome as a modulator of healthy ageing. Nat Rev Gastroenterol Hepatol. 2022 Sep;19(9):565–84.

27. QIAGEN. QIAamp Fast DNA Stool Mini Kit [Internet]. QIAGEN. [cited 2025 Oct 28]. Available from: https://www.qiagen.com/us/products/discovery-and-translational-research/dna-rna-purification/dna-purification/genomic-dna/qiaamp-fast-dna-stool-mini-kit?catno=51604

28. Bolyen E, Rideout JR, Dillon MR, Bokulich NA, Abnet CC, Al-Ghalith GA, et al. Reproducible, interactive, scalable and extensible microbiome data science using QIIME 2. Nat Biotechnol. 2019 Aug;37(8):852–7.

29. Molano L-AG, Vega-Abellaneda S, Manichanh C. GSR-DB: a manually curated and optimized taxonomical database for 16S rRNA amplicon analysis. mSystems. 2024 Feb 20;9(2):e0095023.

30. R Core Team. R: A language and environment for statistical computing. Vienna, Austria: R Foundation for Statistical Computing; 2025.

31. Breiman L. Random Forests. Springer Science and Business Media LLC. 2001;45:5–32.

32. Lefebo BK, Kassa DH, Tarekegn BG. Factors associated with stunting: gut inflammation and child and maternal-related contributors among under-five children in Hawassa City, Sidama Region, Ethiopia. BMC nutrition. 2023;9(1):54.

33. Karlsson O, Kim R, Moloney GM, Hasman A, Subramanian SV. Patterns in child stunting by age: A cross-sectional study of 94 low- and middle-income countries. Matern Child Nutr. 2023 Oct;19(4):e13537.

34. Ekholuenetale M, Okonji OC, Nzoputam CI, Barrow A. Inequalities in the prevalence of stunting, anemia and exclusive breastfeeding among African children. BMC Pediatr. 2022 Jun 9;22(1):333.

35. Thompson AL, Onyango M, Sakala P, Manda J, Berhane E, Selvaggio MP, et al. Are boys more vulnerable to stunting? Examining risk factors, differential sensitivity, and measurement issues in Zambian infants and young children. BMC Public Health. 2024 Nov 29;24(1):3338.

36. Thompson AL. Greater male vulnerability to stunting? Evaluating sex differences in growth, pathways and biocultural mechanisms. Ann Hum Biol. 2021 Sep;48(6):466–73.

37. Hossain MA, Karim MR, Al Mamun ASM, Islam MS, Hossain MG. Logistic regression analysis of mode of delivery influencing nutritional status among under-five children in Bangladesh: survey in rural area of Rajshahi District. International Journal of Statistical Sciences. 2023;23(2):107–16.

38. Carr LE. The role of the infant microbiota and childhood stunting in rural Zimbabwe. 2021;

39. Judijanto L, Budiastutik I, Sugiatmi S, Marlenywati M, Trisnawati E, Kusbandiyah J, et al. Exploring the role of gut Microbiota diversity in early childhood stunting: a multi-center longitudinal study. Evol Stud Imaginative Cult. 2024;8(1):681–90.

40. Zambruni M, Ochoa TJ, Somasunderam A, Cabada MM, Morales ML, Mitreva M, et al. Stunting Is Preceded by Intestinal Mucosal Damage and Microbiome Changes and Is Associated with Systemic Inflammation in a Cohort of Peruvian Infants. Am J Trop Med Hyg. 2019 Nov;101(5):1009–17.

41. Surono IS, Popov I, Verbruggen S, Verhoeven J, Kusumo PD, Venema K. Gut microbiota differences in stunted and normal-lenght children aged 36-45 months in East Nusa Tenggara, Indonesia. PLoS ONE. 2024 Mar 29;19(3):e0299349.

42. Jiang L, Shang M, Yu S, Liu Y, Zhang H, Zhou Y, et al. A high-fiber diet synergizes with Prevotella copri and exacerbates rheumatoid arthritis. Cell Mol Immunol. 2022 Dec;19(12):1414–24.

43. Dinh DM, Ramadass B, Kattula D, Sarkar R, Braunstein P, Tai A, et al. Longitudinal analysis of the intestinal microbiota in persistently stunted young children in south india. PLoS ONE. 2016 May 26;11(5):e0155405.

44. Meiners F, Ortega-Matienzo A, Fuellen G, Barrantes I. Gut microbiome-mediated health effects of fiber and polyphenol-rich dietary interventions. Front Nutr. 2025 Aug 29;12:1647740.

45. Louis P, Flint HJ. Formation of propionate and butyrate by the human colonic microbiota. Environ Microbiol. 2017 Jan;19(1):29–41.

46. Raimondi S, Musmeci E, Candeliere F, Amaretti A, Rossi M. Identification of mucin degraders of the human gut microbiota. Sci Rep. 2021 May 27;11(1):11094.

47. Wu X, Yang H-J, Ryu M-S, Jung S-J, Ha K, Jeong D-Y, et al. Association of Mucin-Degrading Gut Microbiota and Dietary Patterns with Colonic Transit Time in Constipation: A Secondary Analysis of a Randomized Clinical Trial. Nutrients. 2024 Dec 31;17(1).

48. Sokol H, Pigneur B, Watterlot L, Lakhdari O, Bermúdez-Humarán LG, Gratadoux J-J, et al. Faecalibacterium prausnitzii is an anti-inflammatory commensal bacterium identified by gut microbiota analysis of Crohn disease patients. Proc Natl Acad Sci USA. 2008 Oct 28;105(43):16731–6.

49. Ruigrok RAAA, Collij V, Sureda P, Klaassen MAY, Bolte LA, Jansen BH, et al. The composition and metabolic potential of the human small intestinal microbiota within the context of inflammatory bowel disease. J Crohns Colitis. 2021 Aug 2;15(8):1326–38.

50. Louis P, Duncan SH, Sheridan PO, Walker AW, Flint HJ. Microbial lactate utilisation and the stability of the gut microbiome. Gut Microb. 2022;3.

51. Deering KE, Devine A, O’Sullivan TA, Lo J, Boyce MC, Christophersen CT. Characterizing the composition of the pediatric gut microbiome: A systematic review. Nutrients. 2019 Dec 19;12(1).

52. Gatya M, Fibri DLN, Utami T, Suroto DA, Rahayu ES. Gut Microbiota Composition in Undernourished Children Associated with Diet and Sociodemographic Factors: A Case-Control Study in Indonesia. Microorganisms. 2022 Aug 30;10(9).

53. Chibuye M, Mende DR, Spijker R, Simuyandi M, Luchen CC, Bosomprah S, et al. Systematic review of associations between gut microbiome composition and stunting in under-five children. npj Biofilms and Microbiomes. 2024 May 23;10(1):46.

54. Zadock HB, Nzengya DM. Children’s Health Outcomes Associated with Access to Water, Sanitation and Hygiene (WASH): A Systematic Review of Empirical Gaps Related to Diarrhoea in Low-and Middle-Income Countries. African Multidisciplinary Journal of Research. 2025;2(3):24–38.

55. Fusco W, Lorenzo MB, Cintoni M, Porcari S, Rinninella E, Kaitsas F, et al. Short-Chain Fatty-Acid-Producing Bacteria: Key Components of the Human Gut Microbiota. Nutrients. 2023 May 6;15(9).

56. Martin-Gallausiaux C, Marinelli L, Blottière HM, Larraufie P, Lapaque N. SCFA: mechanisms and functional importance in the gut. Proc Nutr Soc. 2021 Feb;80(1):37–49.

57. Schaus SR, Vasconcelos Pereira G, Luis AS, Madlambayan E, Terrapon N, Ostrowski MP, et al. Ruminococcus torques is a keystone degrader of intestinal mucin glycoprotein, releasing oligosaccharides used by Bacteroides thetaiotaomicron. MBio. 2024 Aug 14;15(8):e0003924.

58. Glover JS, Ticer TD, Engevik MA. Characterizing the mucin-degrading capacity of the human gut microbiota. Sci Rep. 2022 May 19;12(1):8456.

59. Precup G, Vodnar D-C. Gut Prevotella as a possible biomarker of diet and its eubiotic versus dysbiotic roles: a comprehensive literature review. Br J Nutr. 2019 Jul 28;122(2):131–40.

60. Toussaint Nguélé A, Mozzicafreddo M, Carrara C, Piersanti A, Salum SS, Ali SM, et al. Interplay Between Helminth Infections, Malnutrition, and Gut Microbiota in Children and Mothers from Pemba, Tanzania: Potential of Microbiota-Directed Interventions. Nutrients. 2024 Nov 24;16(23).

61. Andres SF, Zhang Y, Kuhn M, Scottoline B. Building better barriers: how nutrition and undernutrition impact pediatric intestinal health. Front Immunol. 2023 Jul 21;14:1192936.

62. Laitinen K, Mokkala K. Overall dietary quality relates to gut microbiota diversity and abundance. Int J Mol Sci. 2019 Apr 13;20(8).

63. Maskarinec G, Hullar MAJ, Monroe KR, Shepherd JA, Hunt J, Randolph TW, et al. Fecal Microbial Diversity and Structure Are Associated with Diet Quality in the Multiethnic Cohort Adiposity Phenotype Study. J Nutr. 2019 Sep 1;149(9):1575–84.

64. Ecklu-Mensah G, Gilbert J, Devkota S. Dietary selection pressures and their impact on the gut microbiome. Cell Mol Gastroenterol Hepatol. 2022;13(1):7–18.

65. De Filippo C, Cavalieri D, Di Paola M, Ramazzotti M, Poullet JB, Massart S, et al. Impact of diet in shaping gut microbiota revealed by a comparative study in children from Europe and rural Africa. Proc Natl Acad Sci USA. 2010 Aug 17;107(33):14691–6.

66. Yelverton CA, Killeen SL, Feehily C, Moore RL, Callaghan SL, Geraghty AA, et al. Maternal breastfeeding is associated with offspring microbiome diversity; a secondary analysis of the MicrobeMom randomized control trial. Front Microbiol. 2023 Aug 31;14:1154114.

67. Differding MK, Benjamin-Neelon SE, Hoyo C, Østbye T, Mueller NT. Timing of complementary feeding is associated with gut microbiota diversity and composition and short chain fatty acid concentrations over the first year of life. BMC Microbiol. 2020 Mar 11;20(1):56.

68. Davis JCC, Lewis ZT, Krishnan S, Bernstein RM, Moore SE, Prentice AM, et al. Growth and morbidity of Gambian infants are influenced by maternal milk oligosaccharides and infant gut microbiota. Sci Rep. 2017 Jan 12;7:40466.

69. Verhasselt V, Tellier J, Carsetti R, Tepekule B. Antibodies in breast milk: Pro-bodies designed for healthy newborn development. Immunol Rev. 2024 Nov;328(1):192–204.

70. Ajmal R. Promoting breastfeeding and complementary feeding practices for optimal maternal and child nutrition. Pakistan Journal of Public Health. 2024;14(Special. NI):168–80.

71. Senghor B, Sokhna C, Ruimy R, Lagier J-C. Gut microbiota diversity according to dietary habits and geographical provenance. Human Microbiome Journal. 2018 Feb;7–8:1–9.

72. Ratnayani, Hegar B, Sunardi D, Fadilah F, Gunardi H, Fahmida U, et al. Association of Gut Microbiota Composition with Stunting Incidence in Children under Five in Jakarta Slums. Nutrients. 2024;16(20):3444.

73. Robertson RC, Edens TJ, Carr L, Mutasa K, Gough EK, Evans C, et al. The gut microbiome and early-life growth in a population with high prevalence of stunting. Nat Commun. 2023 Feb 14;14(1):654.

74. Bauza V, Madadi V, Ocharo R, Nguyen TH, Guest JS. Enteric pathogens from water, hands, surface, soil, drainage ditch, and stream exposure points in a low-income neighborhood of Nairobi, Kenya. Sci Total Environ. 2020 Mar 20;709:135344.

75. Rouhani S, Griffin NW, Yori PP, Gehrig JL, Olortegui MP, Salas MS, et al. Diarrhea as a potential cause and consequence of reduced gut microbial diversity among undernourished children in peru. Clin Infect Dis. 2020 Aug 14;71(4):989–99.

76. Li Y, Xia S, Jiang X, Feng C, Gong S, Ma J, et al. Gut microbiota and diarrhea: an updated review. Front Cell Infect Microbiol. 2021 Apr 15;11:625210.

77. Reyman M, van Houten MA, Watson RL, Chu MLJN, Arp K, de Waal WJ, et al. Effects of early-life antibiotics on the developing infant gut microbiome and resistome: a randomized trial. Nat Commun. 2022 Feb 16;13(1):893.

