## Supplementary materials for "Impact of WaSH and dietary practices on age-driven gut microbiome in stunted young children"

**Supplementary Appendices**

Supplementary table 1: Multivariate Logistic Regression (Adjusted Odds Ratios)

| **Term** | **Odds Ratio** | **CI_lower** | **CI_upper** | **P_value** |
| --- | --- | --- | --- | --- |
| (Intercept) | 1.478 | 0.056 | 264.368 | 0.830 |
| Age in months | 1.060 | 0.984 | 1.143 | 0.125 |
| **Gender: male** | **1.998** | **1.173** | **3.440** | **0.011** |
| **Underweight status: Underweight** | **6.335** | **2.306** | **21.635** | **0.000** |
| Biological mother education level: colleges or university | 0.235 | 0.001 | 4.488 | 0.364 |
| Biological mother education level: never attended school | 3.360 | 0.015 | 780.935 | 0.590 |
| Biological mother education level: primary school | 0.513 | 0.003 | 8.452 | 0.676 |
| Biological mother education level: secondary school | 0.390 | 0.003 | 6.355 | 0.546 |
| Hhs toilet sharing: yes | 0.651 | 0.355 | 1.185 | 0.160 |
| Extended breastfeeding: yes | 0.720 | 0.314 | 1.640 | 0.434 |
| Dietary diversity status: poor diversity | 0.639 | 0.199 | 1.967 | 0.436 |
| **Residence by ward: Makorongoni** | **0.266** | **0.074** | **0.883** | **0.030** |
| Residence by ward: Nyalumbu | 0.820 | 0.281 | 2.238 | 0.703 |
| Residence by ward: Nyang’oro | 0.545 | 0.172 | 1.638 | 0.282 |
| Residence by ward: Upendo | 0.678 | 0.224 | 1.926 | 0.471 |
| Child handwashing before food: yes | 1.625 | 0.675 | 3.979 | 0.279 |
| Child handwashing after food: yes | 0.968 | 0.423 | 2.180 | 0.937 |
| Bold indicates statistical significance at p < 0.05. | | | | |

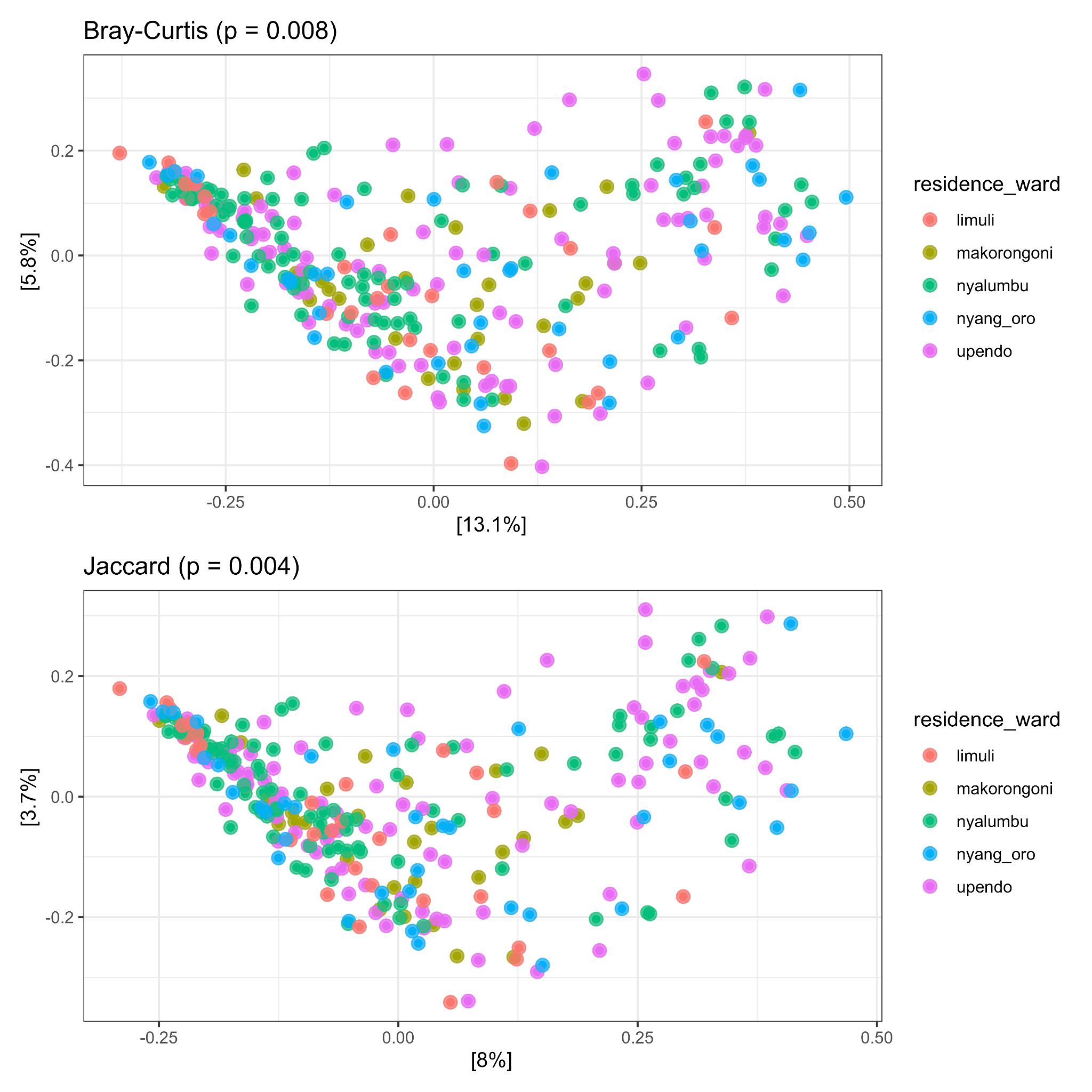

**SF 1**: Beta diversity of gut microbiota by residential ward. Principal Coordinates Analysis (PCoA) plots showing the compositional dissimilarity of gut microbiota across five residential wards (Limuli, Makorongoni, Nyalumbu, Nyang'oro, and Upendo). (Top) Bray-Curtis dissimilarity (PERMANOVA p = 0.008) and (Bottom) Jaccard distance (PERMANOVA p = 0.004). Each dot represents one sample, coloured by residential ward. Statistical significance was assessed using PERMANOVA with 999 permutations.

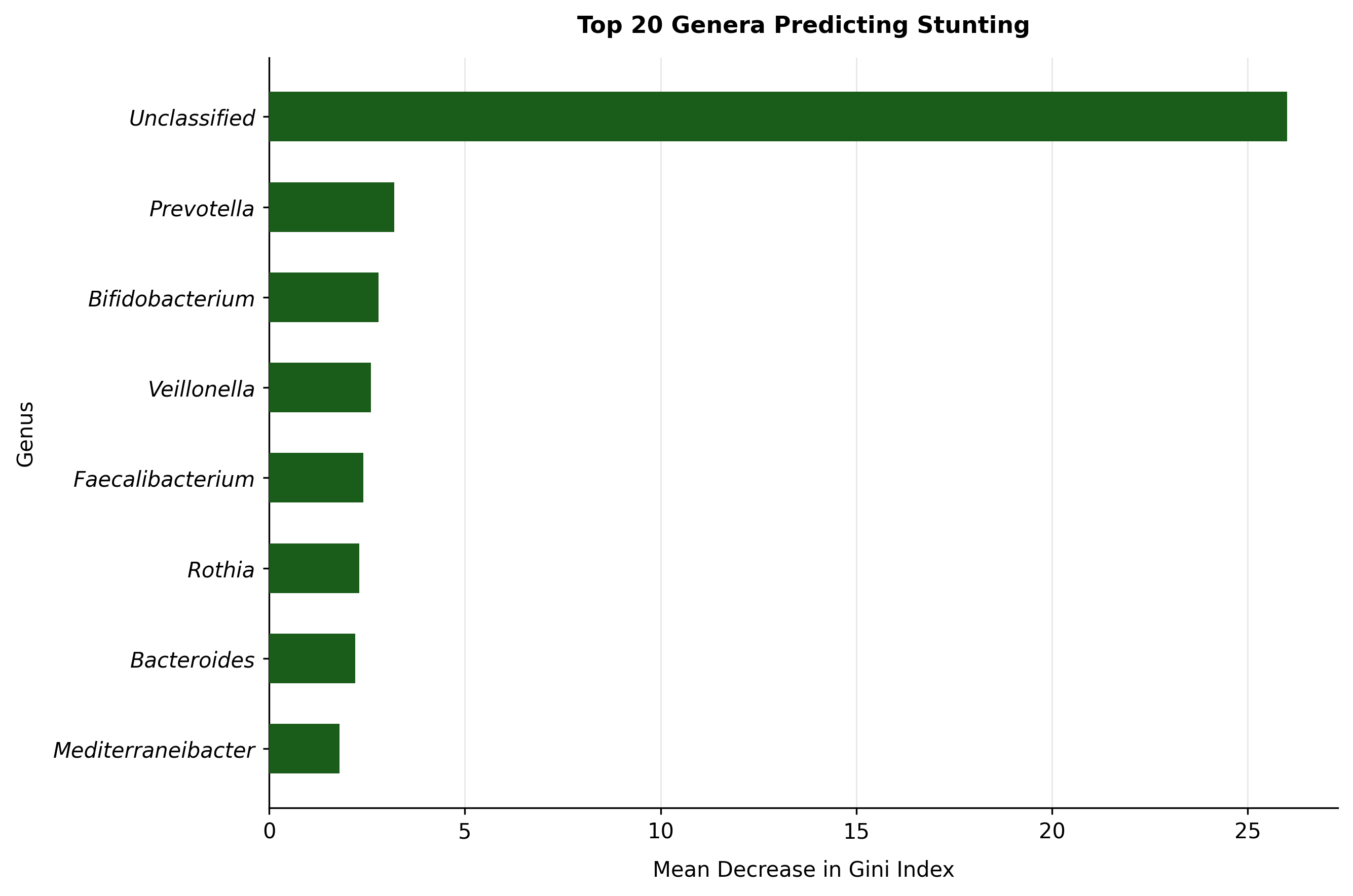

**SF 2**: Random forest model identifying microbial genera as predictors of stunting status. Bar chart showing the mean decrease in Gini index for the top genera selected by a random forest classifier trained to predict stunting status, achieving 87.5% accuracy based on 10-fold cross-validation
